# Nr2f1a maintains atrial *nkx2.5* expression to repress pacemaker identity within venous atrial cardiomyocytes

**DOI:** 10.1101/2022.02.24.481762

**Authors:** Kendall E. Martin, Padmapriyadarshini Ravisankar, Manu E. M. Beerens, Calum A. MacRae, Joshua S. Waxman

## Abstract

Maintenance of cardiomyocyte identity is vital for normal heart development and function. However, our understanding of cardiomyocyte plasticity remains incomplete. Here, we show that sustained expression of the zebrafish transcription factor (TF) Nr2f1a prevents the progressive acquisition of ventricular cardiomyocyte (VC) and pacemaker cardiomyocyte (PC) identities within distinct regions of the atrium. Transcriptomic analysis of isolated atrial cardiomyocytes (ACs) from *nr2f1a* mutant zebrafish embryos showed increased VC marker gene expression and altered expression of core PC regulatory genes, including decreased expression of *nkx2.5*, a critical repressor of PC differentiation. At the arterial pole of the atrium in *nr2f1a* mutants, cardiomyocytes resolve to VC identity within the expanded atrioventricular canal. However, at the venous pole, there is a progressive wave of AC transdifferentiation into PCs across the atrium toward the arterial pole. Restoring Nkx2.5 is sufficient to repress PC identity in *nr2f1a* mutant atria and analysis of chromatin accessibility identified a Nr2f1a-dependent *nkx2.5* enhancer expressed in the atrial myocardium directly adjacent to PCs, supporting that Nr2f1a limits PC differentiation within venous ACs via maintaining *nkx2.5* expression. The Nr2f-dependent maintenance of AC identity within discrete atrial compartments may provide insights into the molecular etiology of concurrent structural congenital heart defects and associated arrhythmias.

## Introduction

The vertebrate heart relies on the coordination of multiple specialized cardiomyocyte populations. Cardiomyocytes of the atrium, ventricle, atrioventricular canal, and pacemaker are each characterized by distinct mechanical, morphological, and electrophysiological properties (Bootman et al., 2006; Brandenburg et al., 2016; Christoffels et al., 2010; Keith & Flack, 1907; Ng et al., 2010; Smyrnias et al., 2010). Pacemaker cardiomyocytes (PCs) of the sinoatrial node (SAN), which is located at the venous pole of the single atrial chamber in zebrafish and the base of the right atrial chamber in mammals, initiate the synchronized wave of contraction that passes through the working atrial cardiomyocytes (ACs), slows at the atrioventricular canal (AVC), and then rapidly activates ventricular cardiomyocytes (VCs) (Arrenberg et al., 2010; Burkhard et al., 2017; Christoffels et al., 2010; Moorman & Christoffels, 2003). Specific gene regulatory programs confer the differentiation and maintenance of these different cardiomyocyte populations within the heart during development (Barth et al., 2005; Ng et al., 2010; Pradhan et al., 2017; Tabibiazar et al., 2003; Targoff et al., 2013; van Weerd & Christoffels, 2016; Xin et al., 2007). Importantly, mutations in genes associated with promoting cardiomyocyte differentiation as well as the maintenance of identity, such as the transcription factors (TFs) Nkx2.5, Tbx5, and Nr2f2, are associated with human congenital heart defects (CHDs), the most common type of congenital malformations found in newborn children (Al Turki et al., 2014; Benjamin et al., 2018; Benson et al., 1999; Cheng et al., 2011; Hoffman & Kaplan, 2002; Loffredo, 2000; Nakamura et al., 2011; Schott et al., 1998; van der Linde et al., 2011). Moreover, structural CHDs are often accompanied by arrhythmias that cause additional complications, morbidity, and death (Bruneau et al., 1999; Ellesøe et al., 2016; Williams & Perry, 2018). In adults, variants in *Nkx2.5* and *Tbx5* are also associated with isolated arrhythmias, such as atrial fibrillation, and improper cardiomyocyte gene expression (Benson et al., 1999; Bruneau et al., 1999; Guo et al., 2016; Jhaveri et al., 2018; Ma et al., 2016; Nakashima et al., 2014; Yu et al., 2014). Thus, the acquisition of an appropriate number and maintenance of identity of each cardiomyocyte population within the developing heart is vital for its normal function throughout life.

The Nr2f (Coup-tf) family of orphan nuclear hormone receptor TFs are conserved regulators of atrial development. In humans, mutations in *NR2F2* have been associated with multiple types of CHDs, most commonly atrioventricular septal defects (AVSDs) (Al Turki et al., 2014; Nakamura et al., 2011; Qiao et al., 2018; Upadia et al., 2018). Consistent with a role in AC differentiation, NR2F1 and NR2F2 are expressed in ACs of the developing hearts of mice and humans (Al Turki et al., 2014; G. Li et al., 2016). *In vitro* studies have shown that both NR2F1 and NR2F2 promote AC differentiation in human embryonic stem cell (ESC)-derived cardiomyocytes (Churko et al., 2018; Devalla et al., 2015). Global Nr2f2 knockout (KO) mice are embryonic lethal with prominent heart defects, including severely hypomorphic atria that lack septa, which is associated with reduced AC differentiation (Pereira et al., 1999). However, at subsequent stages, Nr2f2 appears to maintain atrial identity by directly repressing TFs including *Irx4* and *Hey2,* which promote and maintain the differentiation of VCs. Hence, cardiac-specific conditional KO of *Nr2f2* in mice results in ventricularized atria, while its ectopic expression in VCs is sufficient to promote AC identity (Wu et al., 2013). Although Nr2f2 appears to be the primary regulator of AC differentiation and identity *in vivo* in mammals, we have found that zebrafish Nr2f1a is the functional homolog of mammalian Nr2f2 with respect to heart development (Dohn et al., 2019; Duong et al., 2018). *Nr2f1a* mutant zebrafish develop smaller atria due to reduced AC differentiation at the venous pole and the AVC, which results in an expansion of the AVC into the atrium (Duong et al., 2018). Despite the established requirements for Nr2f TFs in promoting and maintaining atrial differentiation, the mechanisms by which Nr2f TFs function within vertebrate ACs are still not well understood.

Interactions between different cardiomyocyte regulatory programs determine the number and proportions of ACs, VCs, and PCs that initially differentiate and are continuously required to reinforce their identity, which maintains compartmental boundaries within the heart that allow it to function properly. At the venous pole of the heart, a conserved regulatory network of TFs including Isl1, Shox/Shox2, and Tbx3 drive PC differentiation and concurrently repress AC identity. Loss of these TFs results in ectopic expression of working AC markers within the SAN and, in turn, a hypoplastic SAN, while ectopic Tbx3 in in working murine cardiomyocytes is sufficient to confer PC identity and overexpression of Shox2 in murine ESCs during differentiation leads to an upregulation of PC genes (Espinoza-Lewis et al., 2009, 2011; Hoogaars et al., 2007; Ionta et al., 2015; Liang et al., 2015; Liu et al., 2011; Tessadori et al., 2012; van Weerd & Christoffels, 2016). Nkx2.5 performs multiple functions within cardiomyocytes during heart development, consistent with pleiotropic structural and conduction defects found in patients with *NKX2.5* mutations (Benson et al., 1999; Ellesøe et al., 2016; Elliott et al., 2003; Jhaveri et al., 2018; McElhinney et al., 2003; Schott et al., 1998; Xie et al., 2013; Yu et al., 2014). At the venous pole of the heart, it is a critical repressor of the PC differentiation regulatory network within working ACs, as loss of Nkx2.5 results in an expansion of PC marker identity within the atrium of mice and fish (Colombo et al., 2018; Espinoza-Lewis et al., 2011; Nakashima et al., 2014). Within VCs at the arterial pole of the heart, Nkx2.5, as well as the TFs Irx4 and Hey2, maintain ventricular identity and repress the atrial gene program. Loss of these genes in mice and zebrafish results in ectopic expression of AC markers within the ventricle (Bao et al., 1999; Bruneau et al., 2000; Koibuchi & Chin, 2007; Pradhan et al., 2017; Targoff et al., 2013). Moreover, in embryonic zebrafish hearts, AC identity is repressed in VCs by FGF signaling, which functions upstream of Nkx2.5 and by inhibition of BMP (de Pater et al., 2012; Pradhan et al., 2017) . However, in mice, FGF signaling is also necessary for proper ventricular conduction through regulation of connexin 43 (Sakurai et al., 2013). While the repression of AC identity is central to normal development of the ventricle and SAN, we still lack an understanding of mechanisms that maintain AC identity and plasticity within the embryonic vertebrate heart.

Here, we demonstrate that Nr2f1a is required to inhibit the progressive acquisition of VC and PC identity within different regions of the embryonic zebrafish atria. At the arterial pole of the atrium, Nr2f1a represses VC identity, similar to what has been shown in conditional Nr2f2 KO mice. However, we find it is only sensitized cardiomyocytes within the expanded AVC of *nr2f1a* mutants, which co-express AC and VC differentiation markers, that progressively resolve to express only VC markers. Importantly, in venous pole ACs, we identify a requirement for Nr2f1a in repressing PC identity. While the initial PC population differentiates normally in *nr2f1a* mutants despite reduced AC differentiation, there is a subsequent expansion of PC markers and complementary loss of Nkx2.5 from the venous pole into the atrium. Electrophysiological studies demonstrate that the transdifferentiated ACs in *nr2f1a* mutants are adopting conduction system identity. Genetic epistasis using a heat-shock inducible *nkx2.5* transgene showed that overexpressing Nkx2.5 is sufficient to inhibit the transdifferentiation of ACs that occurs in *nr2f1a* mutant hearts, indicating that within ACs Nr2f1a functions upstream of Nkx2.5 in the conserved genetic hierarchy that limits PC differentiation. Chromatin accessibility integrated with our transcriptomic analysis of sorted ACs identified a putative Nr2f1a-dependent *nkx2.5* enhancer that is expressed in a specific subset of venous ACs bordering the PCs in the SAN. Overall, our results demonstrate that maintenance of Nr2f expression is critical for normal vertebrate heart development through sustaining atrial identity in different AC subpopulations, which may provide insights into the etiology of concomitant congenital structural cardiac and conduction defects found in humans.

## Results

### Cardiomyocytes within the AVC progressively resolve to VC identity in *nr2f1a* mutants

Although previous studies have indicated that Nr2f TFs are required to promote atrial differentiation (Duong et al., 2018; Pereira et al., 1999; Wu et al., 2013), we still do not fully understand the effects of Nr2f loss on AC identity. To investigate the consequences of Nr2f loss within the atrium, we performed RNA-seq on captured ACs from wild-type (WT) and *nr2f1a* mutant embryos. Embryos carrying the *Tg(amhc:EGFP)* transgene (R. Zhang et al., 2013) were dissociated at 48 hours post-fertilization (hpf) and fluorescence-activated cell sorting (FACS) was performed to isolate GFP+ ACs (**Figure 1A**). The resulting RNA-seq dataset revealed 2077 upregulated and 1,297 downregulated genes with >2-fold change in the *nr2f1a* mutant ACs relative to WT (**Figure 1 – Source Data 1**). Genes associated with AC differentiation were decreased in the *nr2f1a* mutant atrium, confirming the success of this approach (**Figure 1B; Figure 1 – Source Data 1**). However, we also observed two additional trends in the *nr2f1a* mutant ACs: an increase in the expression of genes associated with VC and arterial pole identity; and misexpression of core genes associated with PC differentiation, with genes shown to promote PC differentiation, such as *tbx3a* and *shox* being increased, while *nkx2.5*, a critical repressor of PC differentiation was decreased (**Figure 1B**). Thus, transcriptional analysis implies that *nr2f1a* mutants have ectopic expression of genes promoting VC and PC differentiation with ACs.

**Figure 1.**
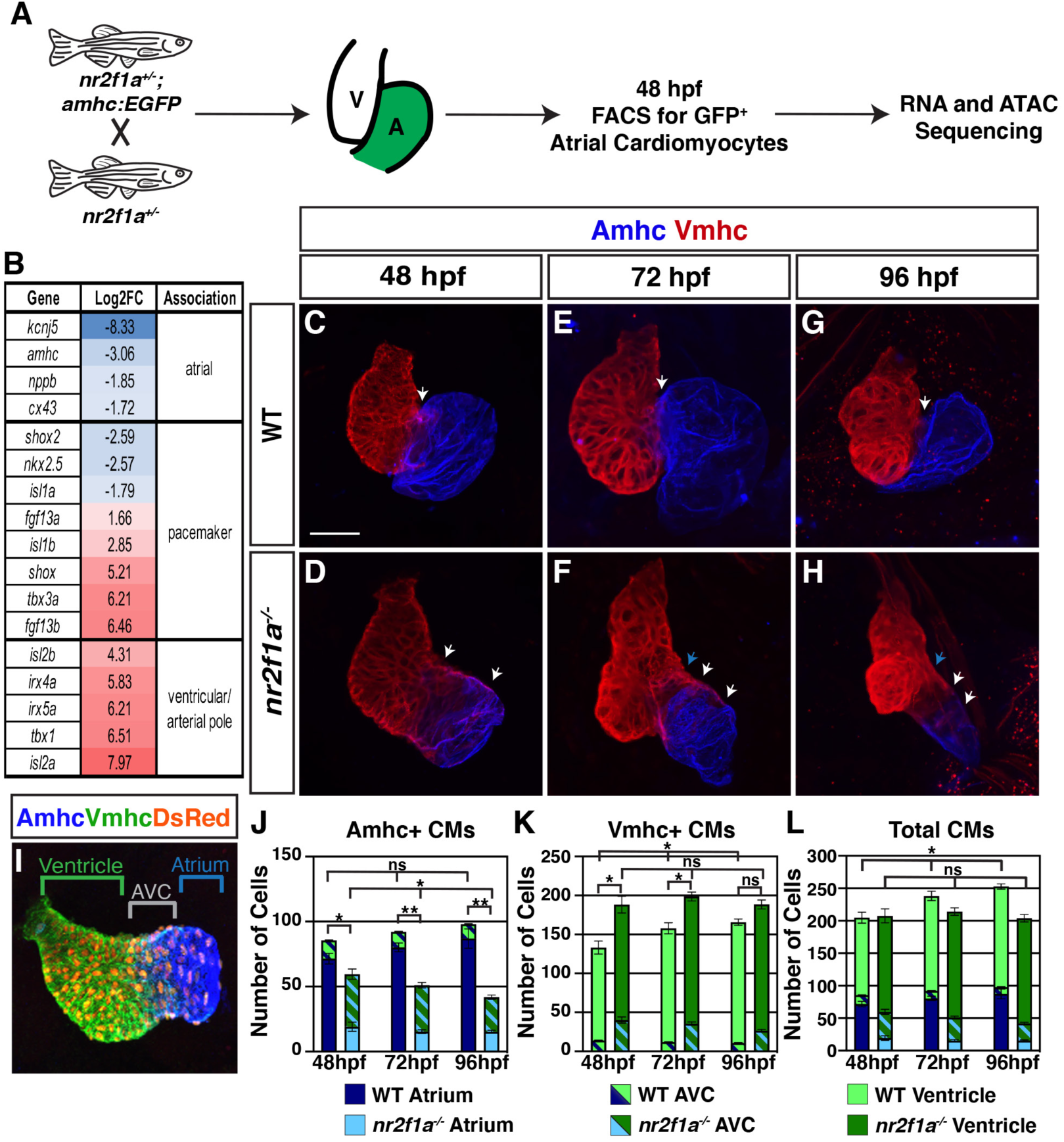
Vmhc expression expands into the AVC of *nr2f1a* mutants. **A**) Schematic for isolation of Acs using *Tg(amhc:EGFP)* transgene for RNA-seq and ATAC-seq at 48 hpf. **B**) Differential expression genes associated with VC/arterial pole, AC, and PC differentiation in *nr2f1a* mutants compared to WT. **C-H**) IHC for Amhc (blue) and Vmhc (red) in *nr2f1a* and WT hearts at 48, 72, and 96 hpf. White arrows indicate region of overlapping Amhc and Vmhc in the AVC. Blue arrows indicate overt arterial position of the AVC of *nr2f1a* mutant atria. 48 hpf: WT (n = 3), *nr2f1a^-/-^* (n = 4); 72 hpf: WT (n = 4), *nr2f1a^-/-^* (n = 6); 96 hpf: WT (n = 7), *nr2f1a^-/-^* (n = 10). **I**) Schematic for cell quantification. Vmhc^+^/Amhc^-^ (green) cardiomyocytes were counted as VCs, Vmhc^+^/Amhc^+^ (green/blue) double-positive cardiomyocytes were counted as AVC, Vmhc^-^/Amhc^+^ (blue) cardiomyocytes were counted as ACs. **J-L)** Quantification of ACs (Amhc^+^-only), AVC cardiomyocytes (Amhc^+^/Vmhc^+^), and VCs (Vmhc^+^-only). Statistics in graphs indicate comparisons between column totals. Comparisons between individual cell populations are shown in Figure 1 - figure supplement 1. Error bars indicate s.e.m. 48 hpf: WT (n = 4), *nr2f1a^-/-^* (n = 5); 72 hpf: WT (n = 5), *nr2f1a^-/-^* (n = 8); 96 hpf: WT (n = 5), *nr2f1a^-/-^* (n = 12). Scale bars indicate 50 μm. Differences between WT and *nr2f1a^-/-^* were analyzed using ANOVA with multiple comparisons. * p = 0.05 – 0.001, ** p < 0.001

Conditional KO of Nr2f2 in murine ACs results in their acquisition of VC identity (Wu et al., 2013). Furthermore, while our transcriptomic analysis here and previous analysis of the ventricular differentiation marker *ventricular myosin heavy chain* (*vmhc*; also called *myh7*) (Duong et al., 2018) also suggests there may be an expansion of VC/arterial pole identity into the atrium of *nr2f1a* mutants, it was still not clear there is a fate switch of ACs to VCs within *nr2f1a* mutant atria at 48 hpf comparable to what was reported in the conditional Nr2f2 KO mice (Wu et al., 2013). To examine AC and VC identity more closely within *nr2f1a* mutant hearts, we performed immunohistochemistry (IHC) for the AC differentiation marker Atrial myosin heavy chain (Amhc; also called Myh6) and Vmhc. In hearts of WT embryos from 48 to 96 hpf, there is minimal overlap of Amhc and Vmhc within the AVC (**Figure 1C,E,G**). However, in the expanded AVC of *nr2f1a* mutants (Duong et al., 2018), we find significant overlap of Amhc and Vmhc by 48 hpf (**Figure 1D**), which gets progressively smaller through 96 hpf, with the Amhc expression domain appearing to recede from the arterial pole of the atrium (**Figure 1F,H**). We corroborated this observation through quantification of the number of cardiomyocytes in the atrium (Amhc^+^-only), AVC (Amhc^+^/Vmhc^+^), and ventricle (Vmhc^+^-only) at 48, 72, and 96 hpf using the *Tg(myl7:DsRed2-NLS)* transgene (Mably et al., 2003) (**Figure 1I**). With respect to ACs, there was a slight increase in total Amhc^+^ (Amhc^+^-only and Amhc^+^/Vmhc^+^) cardiomyocytes in WT hearts over these stages. Conversely, total Amhc^+^ cardiomyocytes were found to decrease from 48 to 96 hpf in *nr2f1a* mutants (**Figure 1J; Figure 1 – figure supplement 1A,B**). With respect to the AVC, the number of Amhc^+^/Vmhc^+^ cardiomyocytes was constant over these stages in WT hearts, while there was a decrease in the number of Amhc^+^/Vmhc^+^ cardiomyocytes in *nr2f1a* mutants (**Figure 1J,K**; **Figure 1 – figure supplement 1B**). With respect to the VCs, there were more total Vmhc^+^ (Vmhc^+^-only and Vmhc^+^/Amhc^+^) cardiomyocytes in *nr2f1a* mutants compared to WT hearts across all timepoints, which reflected the expansion of VC marker identity within the AVC of the *nr2f1a* mutant hearts (**Figure 1K; Figure 1 – figure supplement 1B,C**). Furthermore, the number of Vmhc^+^- only cardiomyocytes progressively increased from 48 to 96 hpf in *nr2f1a* mutant hearts, complementing the loss of Amhc^+^/Vmhc^+^ cardiomyocytes within the AVC (**Figure 1K; Figure 1 – figure supplement 1C**). Interestingly, total cardiomyocyte number increased from 48 to 96 hpf in WT hearts, while the total number of cardiomyocytes in *nr2f1a* mutant hearts remains constant (**Figure 1L**), implying that Nr2f1a loss affects heart growth at these stages. However, we did not observe effects on cell death or proliferation in the hearts of *nr2f1a* mutants (data not shown). Collectively, these results show that the expansion and acquisition of ventricular identity within *nr2f1a* mutant hearts is regionally restricted, with cardiomyocytes comprising the expanded AVC that initially expresses both atrial and ventricular differentiation markers resolving to ventricular identity.

### PC identity progressively expands throughout *nr2f1a* mutant atria

Our results show that not all ACs within the atrium acquire ventricular identity in the absence of Nr2f TFs. Moreover, our transcriptomic data indicated there was also misexpression of core regulatory genes that control PC differentiation in the *nr2f1a* mutant atria (**Figure 1A,B**). Orthologs of genes associated with promoting PC differentiation and function, including *tbx3a*, *shox*, and *fgf13a* (Espinoza-Lewis et al., 2009; Hoogaars et al., 2007; Liu et al., 2011; C. Wang et al., 2011; X. Wang et al., 2017; Yang et al., 2016), were predominantly increased, while *nkx2.5*, a critical repressor of PC identity in mice and zebrafish (Colombo et al., 2018; Hoogaars et al., 2007; Nakashima et al., 2014), was decreased. To determine if there are effects on the PCs in *nr2f1a* mutants, we used the *SqET33-mi59B* enhancer trap line, which for clarity we refer to as *Et(fgf13a:EGFP)*, as it reports *fgf13a* expression and is expressed in PCs (Poon et al., 2016). At 48 hpf, we found no overt difference in *Et(fgf13a:EGFP)* expression between the WT and *nr2f1a* mutant embryos (**Figure 2A,B**), despite the smaller atria in the *nr2f1a* mutants However, while *Et(fgf13a:EGFP)* expression is normally restricted to the venous pole in WT hearts, we found a progressive expansion of *Et(fgf13a:EGFP)* expression from the venous pole throughout the atrium of the *nr2f1a* mutants through 96 hpf (**Figure 2C-F**). Quantification of Amhc^+^ cardiomyocytes co-expressing the *Et(fgf13a:EGFP)* transgene via incorporation of the *Tg(myl7:DsRed2-NLS)* transgene confirmed the concurrent increase in *fgf13a*:EGFP^+^/Amhc^+^ cardiomyocytes and decrease in *fgf13a:*EGFP^-^/Amhc^+^ cardiomyocytes within the *nr2f1a* mutant atria (**Figure 2G,H**), even as the total number of Amhc^+^ cardiomyocytes decrease (**Figure 2I**). Equivalent results were found with IHC for the SAN marker Isl1 (Liang et al., 2015; Sun et al., 2007; Tessadori et al., 2012) (**Figure 2 – figure supplement 1**). Additionally, real-time quantitative PCR (qPCR) on flow-sorted *Tg(myl7:EGFP)*^+^ (Huang et al., 2003) cardiomyocytes (pan-cardiac) at 96 hpf showed an increase in PC differentiation gene expression, as well as *vmhc* expression, and decrease in *amhc* in *nr2f1a* mutant hearts (**Figure 2J,K**). Thus, our data support that venous ACs within *nr2f1a* mutants are acquiring PC gene expression, suggesting they may be progressively transdifferentiating into PCs.

**Figure 2.**
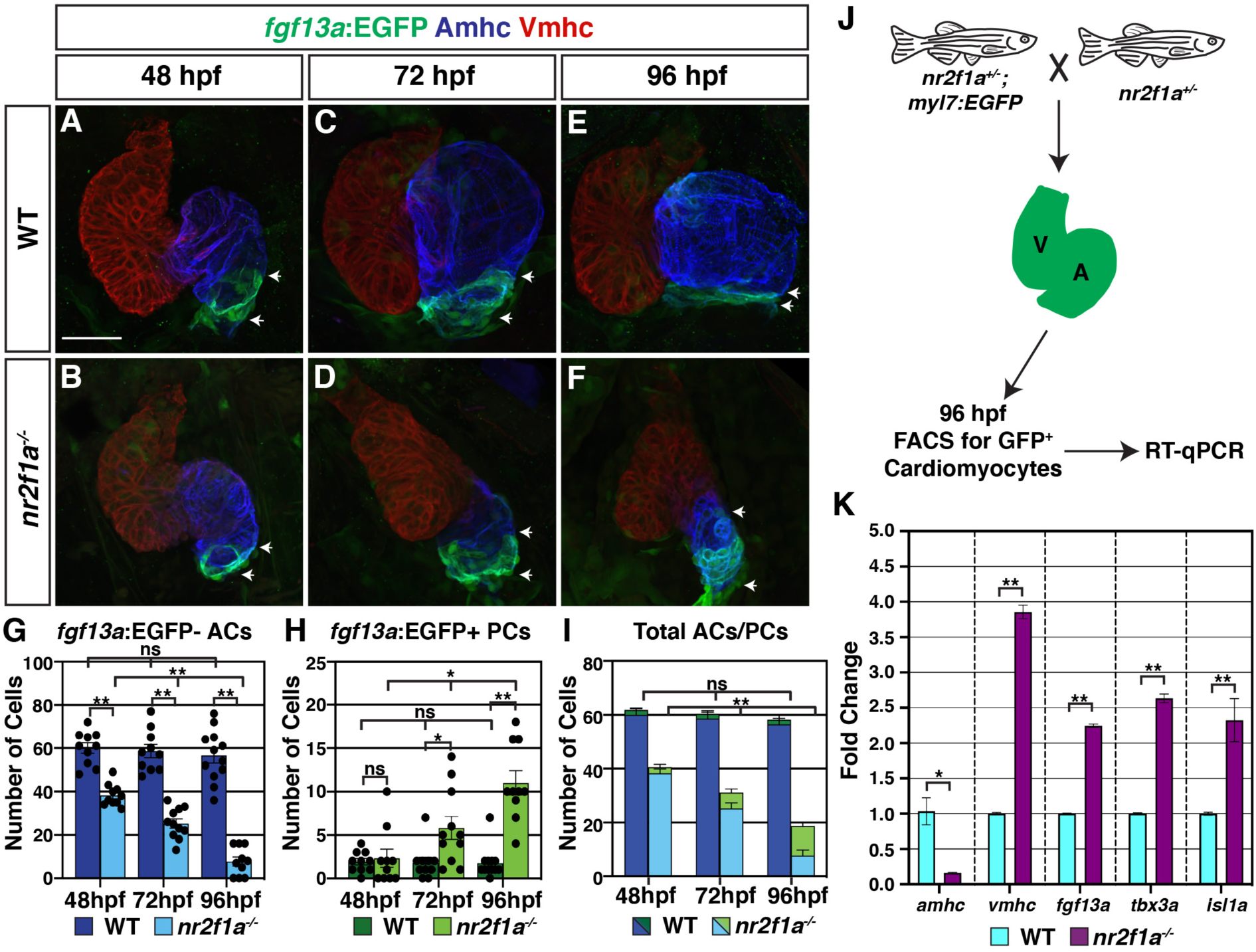
PC identity expands across the atrium in *nr2f1a* mutant embryos. **A-F**) IHC for Amhc (blue), Vmhc (red), and *fgf13a*:EGFP (green). White arrows indicate boundaries of *Et(fgf13a:EGFP)* expression within the atrium. 48 hpf: WT (n = 10), *nr2f1a^-/-^* (n = 12); 72 hpf: WT (n = 10), *nr2f1a^-/-^* (n = 10); 96 hpf: WT (n = 7), *nr2f1a^-/-^* (n = 10). **G-I**) Quantification of Amhc^+^/*fgf13a:*EGFP^+^ cardiomyocytes (PCs) and Amhc^+^/*fgf13a:*EGFP^-^ cardiomyocytes (ACs). Error bars indicate s.e.m. 48 hpf: WT (n = 10), *nr2f1a^-/-^* (n = 10); 72 hpf: WT (n = 10), *nr2f1a^-/-^* (n = 11); 96 hpf: WT (n = 12), *nr2f1a^-/-^* (n = 10). **J**) Schematic for the isolation of cardiomyocytes at 96 hpf using the Tg(*myl7:EGFP)* transgene. **K**) Fold change of marker genes relative to *β-actin* from RT-qPCR on isolated cardiomyocytes of WT and *nr2f1a* mutants at 96 hpf. Scale bars indicate 50 μm. Differences between WT and *nr2f1a^-/-^* were analyzed using ANOVA with multiple comparisons. * p = 0.05 – 0.001, ** p < 0.001

Next, we wanted to determine if the ACs that are acquiring PC gene expression in *nr2f1a* mutants are functionally adopting PC identity. PCs have different electrophysiological properties than working myocardial cells, including slower depolarization and the ability to spontaneously generate action potentials (Christoffels et al., 2010; Lakatta et al., 2010; Mesirca et al., 2015; Schram et al., 2002). Thus, we would predict that if the ACs are adopting conduction/PC-like characteristics the heart rates of *nr2f1a* mutants may be slower than WT embryos. We assessed heart rate using high-speed time lapse imaging (**Figure 3A,B; Figure 3 - Videos 1-6**). At 48 hpf, there was no difference in heart rate between WT and *nr2f1a* mutant hearts (**Figure 3C**). However, as the heart rate of WT embryos increased through 96 hpf, the heart rate of *nr2f1a* mutant hearts failed to increase (**Figure 3C**). To examine the electrophysiologic properties of the ACs that are acquiring PC gene expression in *nr2f1a* mutant hearts, we analyzed myocardial electrophysiology using optical voltage mapping (Colombo et al., 2018; Mosimann et al., 2015; Panáková et al., 2010). The intrinsic bradycardia we observed appeared to be related to effects on the duration and extent of repolarization with evidence of modest hyperpolarization in both heterozygous and homozygous *nr2f1a* hearts, as well as substantial prolongation of repolarization in the homozygous *nr2f1a* hearts as conservatively assessed on the basis of the action potential duration at 20% repolarization (APD20) (**Figure 3D**). Measuring the APD20 within the center of the atria showed that even at 48 hpf the ACs in *nr2f1a* mutant hearts depolarize more slowly compared to WT and heterozygous *nr2f1a* atria (**Figure 3E; Figure 3 – figure supplement 1A**). These effects were not associated with any changes in Phase 4 depolarization (data not shown). There was evidence of significantly slower conduction across the *nr2f1a* mutant atria by 96 hpf (**Figure 3F-L; Figure 3 – figure supplement 1B**), but with no effect on Vmax (**Figure 3 – figure supplement 1C**), consistent with less chamber-like and more conduction system-like intercellular coupling characteristics. Remarkably, there was no evidence of the emergence of the AVC slow conduction zone in the *nr2f1a* mutant hearts that is characteristic of the AV junction and the resultant sequential chamber contraction that is the hallmark of the vertebrate heart (**Figure 3G-L**), supporting that the cardiomyocytes in the AVC of *nr2f1a* mutants have acquired intermediate physiological properties with the VCs. Collectively, our data support that *nr2f1a* mutant ACs are adopting a functional identity that exhibits slowed repolarization and reduced intercellular coupling, closer to the physiology of the conduction system, and paralleling the PC-associated gene expression changes we observe within the *nr2f1a* mutant atria.

**Figure 3.**
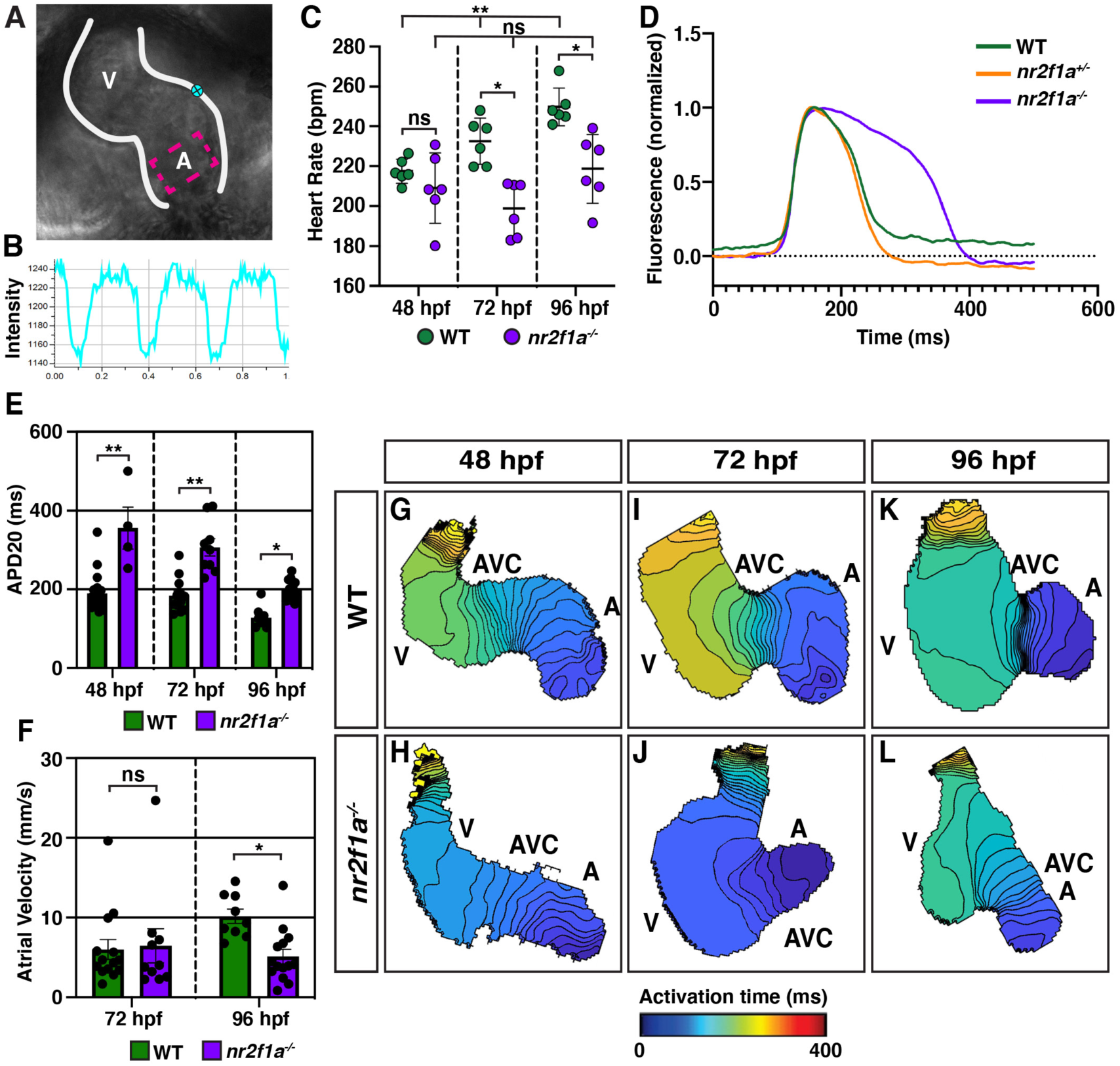
ACs in *nr2f1a* mutants function as PCs. **A**) Schematic of point placements to measure heart rate (cyan) and APD20 (magenta). **B**) Representative kymograph used to analyze heart rate in WT and *nr2f1a* mutant embryos. **C**) Quantification of heart rate in WT and *nr2f1a* mutant embryos. Error bars indicate standard deviation. 48 hpf: WT (n = 6), *nr2f1a^-/-^* (n = 6); 72 hpf: WT (n = 6), *nr2f1a^-/-^* (n = 6); 96 hpf: WT (n = 6), *nr2f1a^-/-^* (n = 6). **D)** Representative atrial action potentials of WT, *nr2f1a^+/-^*, and *nr2f1a^-/-^* embryos at 96 hpf. **E)** APD20 in WT and *nr2f1a^-/-^* hearts. Error bars indicate s.e.m. 48hpf: WT (n = 21), *nr2f1a^-/-^* (n = 4); 72 hpf: WT (n = 13), *nr2f1a^-/-^* (n = 9); 96 hpf: WT (n = 9), *nr2f1a^-/-^* (n = 14). **F)** Atrial Velocity in WT and *nr2f1a^-/-^* hearts. Error bars indicate s.e.m. 72 hpf: WT (n = 13), *nr2f1a^-/-^* (n = 9); 96 hpf: WT (n = 9), *nr2f1a^-/-^* (n = 14). **G-L**) Representative isochronal maps illustrating the positions of the depolarizing wave front in 5 ms intervals for WT and *nr2f1a^-/-^* embryos at 48, 72, and 96 hpf. 48hpf: WT (n=20), *nr2f1a^-/-^* (n=6) 72hpf: WT (n=12), *nr2f1a^-/-^* (n=7) 96hpf: WT (n=11), *nr2f1a^-/-^* (n=14). Differences between WT and *nr2f1a^-/-^* analyzed using ANOVA with multiple comparisons. * p = 0.05 – 0.001, ** p < 0.001

### Nr2f1a represses SAN identity upstream of Nkx2.5

We next wanted to understand how Nr2f1a is regulating the PC gene regulatory network within the atria. We prioritized examining Nkx2.5 in *nr2f1a* mutant hearts, as Nkx2.5 loss in mice and zebrafish results in an expansion of PCs within the atrium (Colombo et al., 2018; Espinoza-Lewis et al., 2011; Nakashima et al., 2014) and its expression was reduced in the bulk RNA-seq data of *nr2f1a* mutant ACs (**Figure 1B**). IHC for Nkx2.5 and the *Et(fgf13a:EGFP)* reporter showed that in WT hearts at 48 through 96 hpf Nkx2.5 is expressed throughout the atrial myocardium but excluded from the PCs at the venous pole (**Figure 4A-C”**). At 48 hpf in *nr2f1a* mutants, Nkx2.5 shows a similar complementary pattern with the *Et(fgf13a:EGFP)* reporter despite the smaller atria (**Figure 4D**). As the *Et(fgf13a:EGFP)* reporter expands into the atria through 96 hpf in *nr2f1a* mutants, Nkx2.5 recedes from the venous pole and maintains the complementary expression pattern (**Figure 4E-F”**). Equivalent results were obtained using the *TgBAC(−36Nkx2.5:ZsYellow)* transgene (Zhou et al., 2011) (**Figure 4 – figure supplement 1**). Thus, a progressive loss of Nkx2.5 within the ACs correlates with and abuts the expansion of PC identity in *nr2f1a* mutant atria.

**Figure 4.**
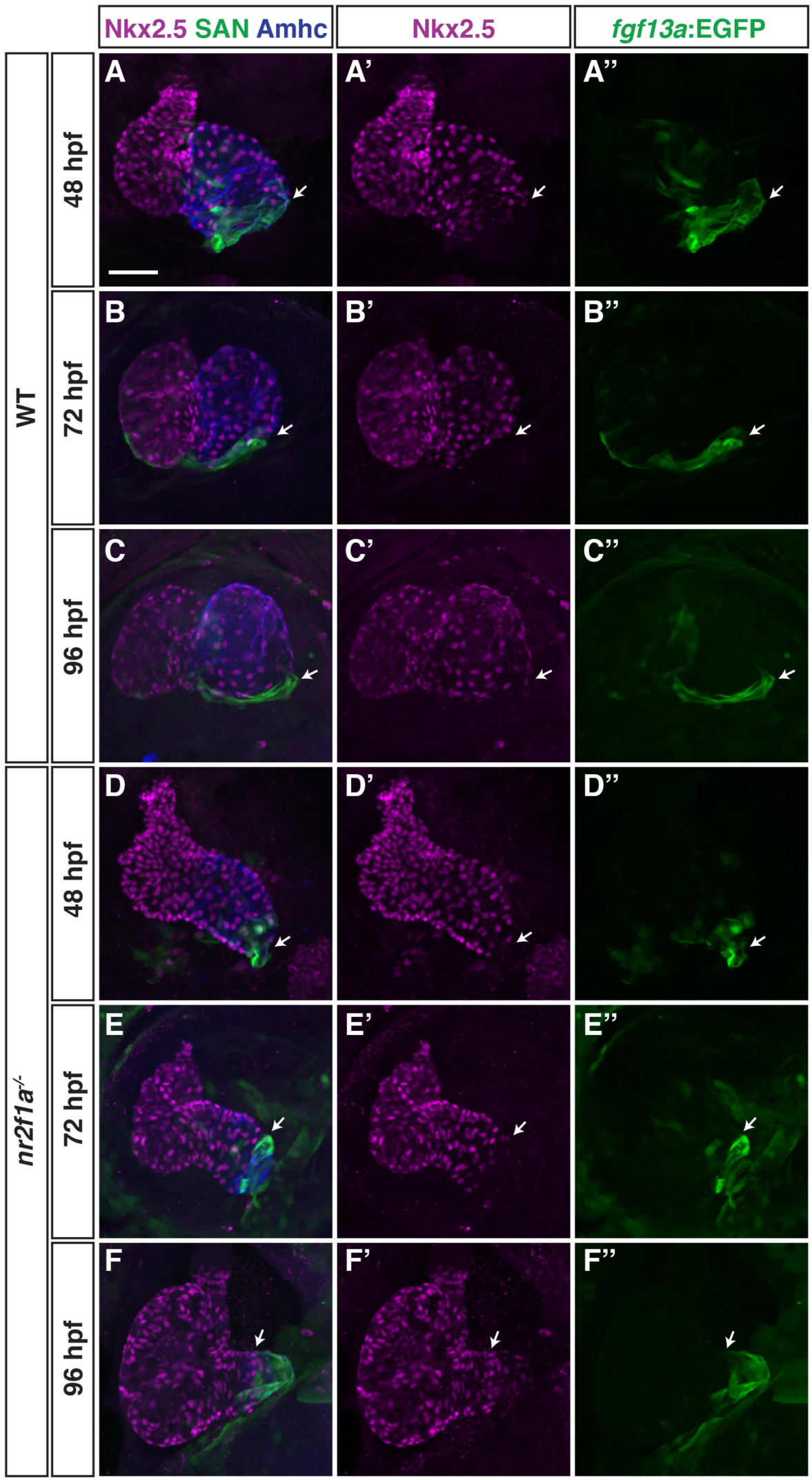
Nkx2.5 expression recedes from venous pole in *nr2f1a* mutant atria. **A-F”**) IHC for Nkx2.5 (purple), Amhc (blue), and *fgf13a:*EGFP (green) in WT and *nr2f1a* mutant embryos from 48 to 96 hpf. White arrows indicate border of *fgf13a:*EGFP^+^ and Nkx2.5^+^ cardiomyocytes. Number of embryos examined at 48 hpf: WT (n = 7), *nr2f1a^-/-^* (n = 6); 72 hpf: WT (n = 7), *nr2f1a^-/-^* (n = 10); 96 hpf: WT (n = 14), *nr2f1a^-/-^* (n = 23). Scale bars indicate 50 μm.

To determine if restoring Nkx2.5 in *nr2f1a* mutants can repress the transdifferentation of ACs to PCs, we used the *Tg(hsp70l:nkx2.5-EGFP)* transgene (George et al., 2015). *Nr2f1a^+/-^* fish that carry the *Tg(myl7:DsRed2-NLS) and Et(fgf13a:EGFP)* transgenes were crossed to *nr2f1a^+/-^* fish carrying the *Tg(hsp70l:nkx2.5-EGFP)* transgene (**Figure 5A**). The resulting embryos were heat-shocked at 20 hpf, as a single heat shock at ∼20 hpf using this transgene is sufficient to rescue *nkx2.5* mutants to adulthood (George et al., 2015). Following heat-shock, the embryos were sorted based on the presence of Nkx2.5-EGFP and allowed to develop until 48 and 96 hpf, at which point the number of PCs were quantified (**Figure 5A**). At 48 hpf, there was a modest decrease in the number of PCs in Nkx2.5-EGFP^+^ WT hearts compared to the Nkx2.5-EGFP^-^ WT siblings (**Figure 5B,C,J,K**). However, Nkx2.5-EGFP^+^ *nr2f1a* mutants had a complete absence of PCs at the venous pole (**Figure 5F,G,J,K**), suggesting that Nkx2.5 is sufficient to repress PC differentiation in *nr2f1a* mutants and, further, that *nr2f1a* mutant ACs may be sensitized to increased Nkx2.5 expression. At 96 hpf, both Nkx2.5-EGFP^+^ WT and *nr2f1a* mutant hearts had an increased number of PCs compared to those at 48 hpf, yet Nkx2.5-EGFP^+^ *nr2f1a* mutants still had significantly fewer PCs than mutants lacking the transgene (**Figure 5D,E,H-K**). We found that heat-shock induction of Nkx2.5-EGFP at 40 hpf could also repress the expansion of PC identity in *nr2f1a* mutant atria (**Figure 5 – figure supplement 1A-C,F,G,J,K**). However, the rescue was not as effective or prolonged as the heat-shock induction of Nkx2.5-EGFP at 20 hpf (**Figure 5 – figure supplement 1D,E,H-K**). Overall, these data suggest that Nr2f1a prevents the expansion of PC identity within the venous pole of the atrium through promoting or maintaining Nkx2.5 in ACs.

**Figure 5.**
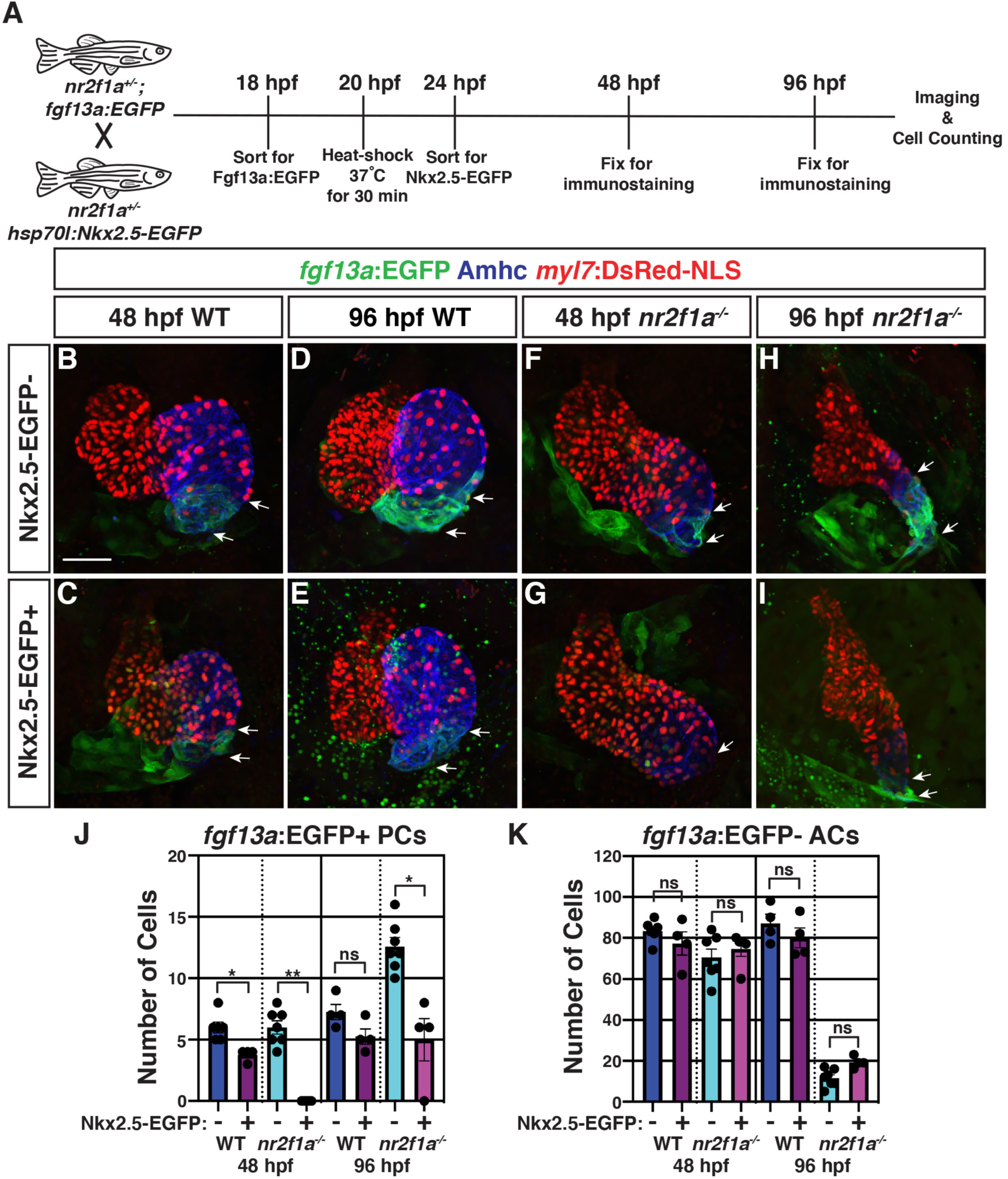
Nkx2.5 induction at 20 hpf represses PC expansion in *nr2f1a* mutant atria. **A**) Schematic of timeline for heat-shock rescue experiments. **B-I**) Representative images of IHC for *myl7:DsRed2-NLS* (red), Amhc (blue), and *fgf13a:*EGFP (green) used to quantify ACs and PCs from WT and *nr2f1a* mutant embryos with and without Nkx2.5-EGFP. White arrows indicate boundaries of *Tg(fgf13a:EGFP)* expression. **J-K**) Quantification of *myl7:*DsRed2-NLS^+^/Amhc^+^/*fgf13a:*EGFP^+^ (PCs) and *myl7:*DsRed2-NLS^+^/Amhc^+^/*fgf13a:*EGFP^-^ (ACs) with and without Nkx2.5-EGFP induction. Error bars indicate s.e.m. 48 hpf WT: Nkx2.5-EGFP^-^ (n = 6), Nkx2.5-EGFP^+^ (n = 4); 48 hpf *nr2f1a^-/-^*: Nkx2.5-EGFP^-^ (n = 7), Nkx2.5-EGFP^+^ (n = 5); 96 hpf WT: Nkx2.5-EGFP^-^ (n = 4), Nkx2.5-EGFP^+^ (n = 4); 96 hpf *nr2f1a^-/-^*: Nkx2.5-EGFP^-^ (n = 7), Nkx2.5-EGFP^+^ (n = 4). Scale bars indicate 50 μm. Differences between WT and *nr2f1a^-/-^* were analyzed using ANOVA with multiple comparisons. * p = 0.05 – 0.001, ** p < 0.001

### A putative *nkx2.5* enhancer is expressed within the venous pole of the atrium

To identify cis-regulatory elements that may regulate *nkx2.5* expression and be affected by Nr2f1a loss, we assayed chromatin accessibility by performing ATAC-seq on flow-sorted ACs from 48 hpf embryos, similar to what is described above (**Figure 1A**). Changes in called peaks of WT and *nr2f1a* mutant ACs were scored relative to the nearest differentially expressed genes (**Figure 1 – Source Data 1**). HOMER was used to determine enrichment for canonical Nr2f TF binding sites on the putative enhancer elements. One of the peaks that was revealed to close in *nr2f1a* mutant atria and contain a Nr2f binding site was ∼55kb upstream of the *nkx2.5* transcriptional start site (**Figure 6A**). This putative enhancer region was placed upstream of the basal *E1b* promoter to drive GFP expression (**Figure 6B**) using the previously reported vector (Birnbaum et al., 2012), and transgenic lines were generated using Tol2 (Kwan et al., 2007). Remarkably, stable transgenic lines showed that this putative enhancer is expressed in a group of cells at the venous pole of the atrium (**Figure 6C**), which are directly adjacent to Isl1^+^ PCs (**Figure 6D,D’**). The *Tg(−55nkx2.5:EGFP)* was crossed into *nr2f1a* carriers to determine if the expression of this putative *nkx2.5*-enhancer is affected in *nr2f1a* mutants. The number of atria with *Tg(−55nkx2.5:EGFP)* expression was significantly decreased in *nr2f1a* mutants compared to WT (**Figure 6C,E,F**). Together, these data suggest that a putative *nkx2.5* enhancer promotes expression adjacent to PCs and requires Nr2f1a within the atria.

**Figure 6.**
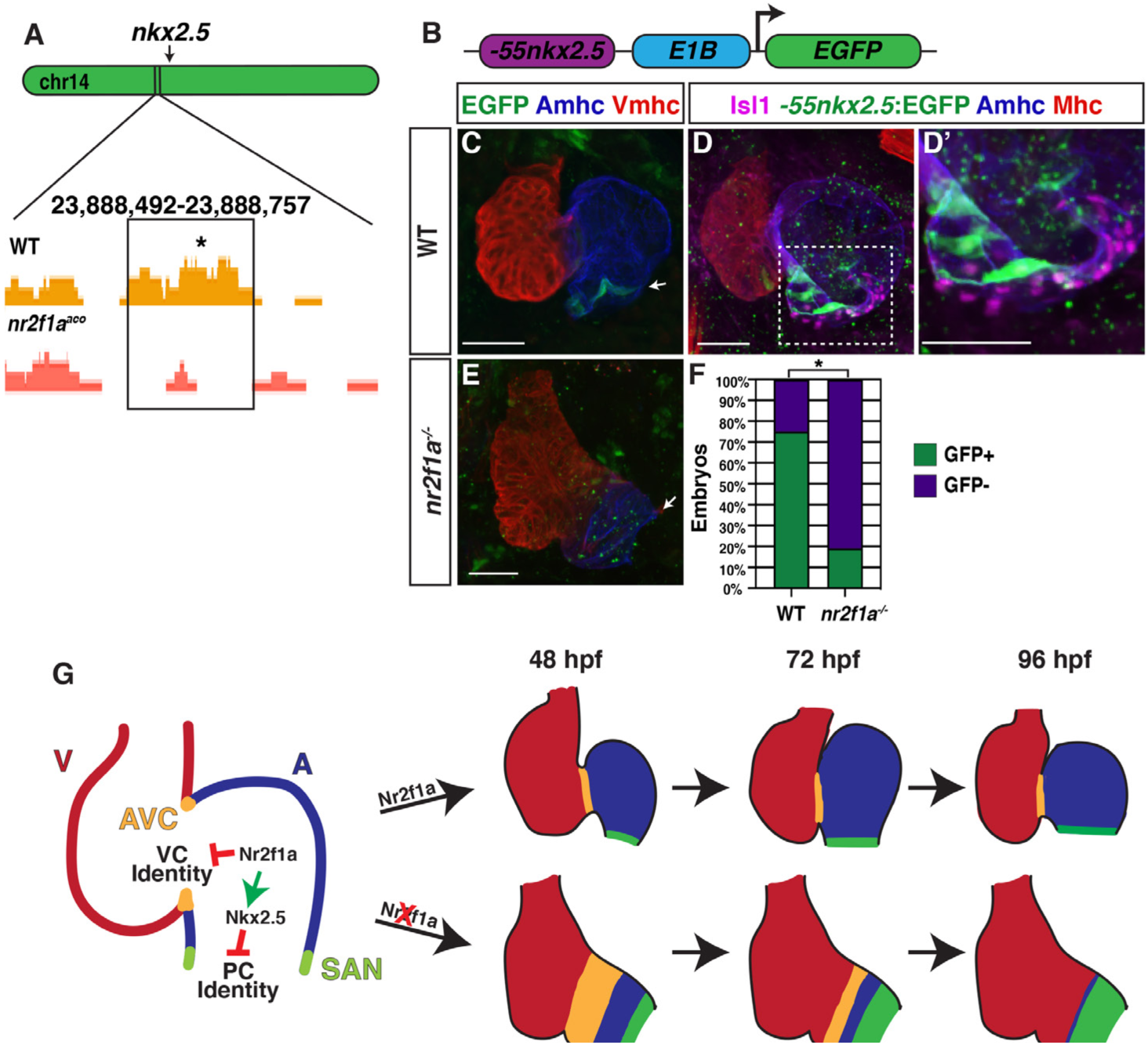
A putative *nkx2.5* enhancer is expressed in ACs adjacent to the SAN. **A**) Comparison of ATAC-seq peaks from WT and *nr2f1a* mutant ACs at 48 hpf, ∼55kb upstream of *nkx2.5*. Asterisk denotes canonical Nr2f binding site. **B**) Schematic of -55kb *nkx2.5*-enhancer reporter construct. **C**) IHC for Vmhc (red), Amhc (blue) and the *Tg(−55nkx2.5:EGFP)* enhancer (green) at 72 hpf. White arrow indicates EGFP expression. **D-D’**) Staining for sarcomeric myosin heavy chain (Mhc: pan-cardiac - red), Amhc (blue), Isl1 (purple), and *-55nkx2.5:EGFP* (green) shows enhancer is expressed in the atrial myocardium directly adjacent to the SAN. **E**) *nr2f1a* mutant at 72 hpf stained for Vmhc (red), Amhc (blue), and *-55nkx2.5:EGFP* (green), cardiac expression of enhancer is lost in *nr2f1a* mutants. White arrow indicates venous pole devoid of *-55nkx2.5:EGFP* expression. **F**) Comparison of percentage of WT and *nr2f1a* embryos with *-55nkx2.5:EGFP* expression in the heart at 72 hpf WT (n = 16); *nr2f1a^-/-^* (n = 16), using Fisher’s exact test. * p < 0.05. Scale bars indicate 50 μm. **G**) Model summarizing the progressive requirements of Nr2f1a in maintaining AC identity within the arterial and venous poles of the heart.

## Discussion

Vertebrate embryonic cardiomyocytes maintain a high degree of plasticity, with their identity being continually reinforced even after they have overtly differentiated (Barth et al., 2005; Ng et al., 2010; Pradhan et al., 2017; Tabibiazar et al., 2003; Targoff et al., 2013; van Weerd & Christoffels, 2016; Xin et al., 2007). Here, our data have advanced our understanding of the requirements of Nr2f TFs within the vertebrate heart and show that they are required to maintain AC identity while concurrently repressing both VC and PC identity in different regions of the atrium (**Figure 6G**). In mice, Nr2f2 maintains AC identity by repressing the expression of TFs that promote a ventricular identity program, including *Irx4* and *Hey2* (Wu et al., 2013). Moreover, murine Nr2f2 appears to be required over a relatively long developmental period after ACs express differentiation markers to prevent the acquisition of VC identity within the atrium. Specifically, loss of Nr2f2 by e9.5 using the *Myh6-Cre* or by e12.5 using the inducible *Myh6-MerCreMer* both result in the ectopic expression of VC differentiation genes within ACs (Wu et al., 2013). Conversely, ectopic Nr2f2 expression in VCs is sufficient to induce AC gene expression (Wu et al., 2013). Although our previous work had shown that zebrafish Nr2f1a promotes atrial differentiation (Duong et al., 2018), it was still unclear if ACs in zebrafish *nr2f1a* mutants acquired VC identity, as shown for *Nr2f2* loss in mice. We now show that there is acquisition of VC identity within the *nr2f1a* mutant atria. However, with the murine conditional Nr2f2 KO approaches, the two atria were formed, but enlarged and appeared to express VC markers throughout the atria (Wu et al., 2013). In contrast to what was shown in the conditional KO mice, the acquisition of VC identity in zebrafish *nr2f1a* mutants was limited to the population of less differentiated cardiomyocytes that comprise the expanded AVC, which extends into the smaller single atria. Moreover, this population of cardiomyocytes within the *nr2f1a* mutant AVC initially express both AC and VC differentiation markers and then progressively lose expression of AC differentiation markers by resolving to VC identity in an arterial-to-venous direction. We presently cannot rule out that these overt differences in effects on the ACs are due to species-specific requirements of Nr2f TFs within mice and zebrafish or technical differences from the global zebrafish *nr2f1a* mutants vs the conditional Nr2f2 approaches. Nevertheless, taken together, the trends reinforce that there are conserved requirements for vertebrate Nr2f TFs in repressing VC identity within ACs. However, our results support that not all ACs have the potential to transdifferentiate into VCs and that this transformation occurs in a sensitized population.

Our data also support a new paradigm whereby Nr2f TFs are required to limit the acquisition of PC identity within more venous ACs. Work predominantly in mice has revealed a transcriptional regulatory network that promotes the differentiation of PCs within the SAN at the venous pole of the right atrium (Burkhard et al., 2017; Christoffels et al., 2010; van Weerd & Christoffels, 2016). Tbx3, Isl1, and Shox/Shox2 TFs are part of a core regulatory network that promotes PC differentiation within venous ACs, while Nkx2.5 represses PC identity in working ACs (Blaschke et al., 2007; Espinoza-Lewis et al., 2009, 2011; Hoogaars et al., 2007; Liang et al., 2015; Liu et al., 2011; Mommersteeg et al., 2007; Nakashima et al., 2014; Sun et al., 2007; Weinberger et al., 2012; Wiese et al., 2009). Although the characterization of PC differentiation has been less extensive in zebrafish, functional analysis of Isl1, Shox/Shox2, and Nkx2.5 support that this regulatory network is fundamentally conserved (Blaschke et al., 2007; Colombo et al., 2018; de Pater et al., 2009; Hoffmann et al., 2013; Tessadori et al., 2012) . Our data show that in *nr2f1a* mutants there is a progressive expansion of conduction system identity from the venous pole toward the arterial pole of the atrium, supporting a role for Nr2f TFs in regulating this conserved network within ACs. The progressive expansion, where the number of PCs is initially unchanged in *nr2f1a* mutant atria despite being smaller, contrasts some with *Nkx2.5* mutant mice and zebrafish (Colombo et al., 2018; Mommersteeg et al., 2007; Nakashima et al., 2014), which show an initial expansion of PC markers and ectopic differentiation at earlier stages. Although the studies of *Nr2f2* conditional KO mice did not examine PC and SAN differentiation, manual interrogation of their data, which was from whole hearts, does show they have increased expression of Tbx3 and Fgf13 (Wu et al., 2013), implying that this requirement in repressing PC identity is also conserved within the vertebrate atrium. However, determining if murine Nr2f2 possesses similar requirements in repressing PC identity at the venous pole of the right atrium in mammals will require additional analysis.

Our data support that one mechanism by which Nr2f1a intersects with the PC regulatory network and represses the progressive acquisition of PC identity within ACs is through maintaining atrial Nkx2.5 expression. While induction of Nkx2.5 can repress PC identity within *nr2f1a* mutant atria, we note that Nkx2.5 induction initially completely represses PC differentiation, implying these ACs are actually sensitized to Nkx2.5 in *nr2f1a* mutants; however, Nkx2.5 induction does not produce a permanent repression of the PC expansion, as the PC markers following Nkx2.5 induction are expanded by 96 hpf, though less than in *nr2f1a* mutant atria. While one interpretation could be that Nkx2.5 needs to be maintained within the ACs over a longer period than provided by the heat-shock to prevent PC expansion, this would seem unlikely as *nkx2.5* mutants can be rescued to adulthood following a single heat-shock at the 20s stage. Therefore, we postulate that Nr2f1a may also concurrently repress the atrial expression of TFs within the core PC regulatory network that promotes PC differentiation, which warrants further investigation. Furthermore, it is interesting that the putative *nkx2.5* enhancer begins expression at later stages (∼72 hpf), correlating with the PC expansion and implying the enhancer may be activated to reinforce *nkx2.5* expression at this border. Additionally, we did not observe overlap of VC and PC markers within the *nr2f1a* mutant atria. As Nr2f1a appears to be expressed equivalently throughout the zebrafish atria (Duong et al., 2018) (**Figure 6 – figure supplement 1**), our data indicate that other factors or signals within the atria must work with Nr2f1a to mediate its regional requirements preventing the progressive acquisition of VC or PC identity.

In considering the implications of our data, it is interesting that in humans, congenital malformations, including AVSDs, are associated with increased incidence of arrhythmias (Williams & Perry, 2018). Moreover, the arrhythmias can arise from complications secondary to the structural defects or pleiotropic requirements in the development of the myocardium and the conduction system (Bruneau et al., 1999; Ellesøe et al., 2016; Williams & Perry, 2018). A prime example of the latter is mutations in *NKX2.5*, which are associated with numerous congenital heart defects, including atrial and ventricular septal defects, and conduction defects, consistent with its requirements in working cardiomyocyte and PC differentiation shown in experimental models (Benson et al., 1999; Ellesøe et al., 2016; Elliott et al., 2003; Jhaveri et al., 2018; McElhinney et al., 2003; Schott et al., 1998; Xie et al., 2013; Yu et al., 2014). Similar to *NKX2.5*, *NR2F2* mutations in humans have now been associated with myriad CHDs, including atrial septal defects (ASDs) and AVSDs (Al Turki et al., 2014; Nakamura et al., 2011; Poot et al., 2007; Qiao et al., 2018; Upadia et al., 2018). However, it is presently unclear if there are also accompanying arrhythmias or conduction defects in these patients. Given our results, we speculate that further evaluation of patients with *NR2F2* mutations will show that they have or are at higher risk to develop arrhythmias.

In conclusion, mutations in genes, such as *NR2F2*, that are required for the maintenance of cardiomyocyte identity have been associated with multiple types of CHDs in humans (Al Turki et al., 2014; Benjamin et al., 2018; Benson et al., 1999; Cheng et al., 2011; Hoffman & Kaplan, 2002; Loffredo, 2000; Nakamura et al., 2011; Schott et al., 1998; van der Linde et al., 2011). Hence, elucidating the transcriptional mechanisms that maintain cardiomyocyte identity and regulate cardiomyocyte plasticity may provide insights into the etiology of CHDs, as well as help refine stem cell-derived tissue engineering and regenerative strategies. Our results have delineated novel requirements for Nr2f TFs in maintaining atrial identity, which may help provide a framework for our understanding of the molecular etiology of concurrent structural malformations and conduction defects in patients.

## Materials and Methods

### Ethics statement

All zebrafish husbandry and experiments were performed in accordance with protocols approved by the Cincinnati Children’s Hospital Medical Center Institutional Animal Care and Use Committee (IACUC).

### Zebrafish lines used

Adult zebrafish were raised and maintained under standard laboratory conditions (Westerfield, 2000). Transgenic lines used: *Tg(−5.1myl7:DsRed2-NLS)^f2^* (Mably et al., 2003), *SqET33-mi59B* (Poon et al., 2016), *TgBAC(−36nkx2.5:ZsYellow)^fb7^* (Zhou et al., 2011), *Tg(myh6:EGFP)^s958^* (R. Zhang et al., 2013), *Tg(hsp70l:nkx2.5-EGFP)^fcu1^* (George et al., 2015), *Tg(myl7:EGFP)^f1^* (Huang et al., 2003). Two *nr2f1a* mutant alleles were used: *nr2f1a^ci1009^*, which was reported previously (Duong et al., 2018), and *nr2f1a^ci1017^*, which was identified from an ENU screen. Additional details of this allele will be reported elsewhere (manuscript in preparation).

### Generation of transgenic Nkx2.5 enhancer line

The putative *nkx2.5* enhancer transgenic line was generated using standard Tol2/Gateway methods (Kwan et al., 2007). The enhancer region at ∼-55kb upstream of *nkx2.5* was amplified with PCR of genomic DNA and cloned into the pDONR221 middle entry vector. Subsequently, it was transferred into the *E1b-GFP-Tol2* Gateway destination vector (Addgene plasmid #37846, Birnbaum et al., 2012). To generate transgenic lines, one cell embryos were co-injected with 25 pg of the *-55nkx2.5:EGFP* plasmid and 25 pg Tol2 mRNA (Kwan et al., 2007). Injected embryos were raised to adulthood and transgenic founders were identified via outcrossing to wild-type fish. Multiple founders with overtly equivalent GFP expression within the atrium were identified. The line that was used in this study has been designated *Tg(−55nkx2.5:EGFP)^ci1015^*.

### Immunohistochemistry (IHC) and cell quantification

IHC was performed as previously described (Waxman et al., 2008). Embryos were fixed in 1% formaldehyde in PBS for 1 hour at room temperature and then washed 1X in PBS, 2X in 0.1% saponin/1X PBS followed by blocking in saponin blocking solution (0.1% saponin, 10% sheep serum, 1X PBS, 2mg/mL BSA) for 1 hour. Primary antibodies were diluted in saponin blocking solution and applied to embryos overnight at 4⁰C. Embryos were washed with 0.1% saponin/1X PBS multiple times before being incubated with secondary antibodies diluted in saponin blocking solution for 2 hours at room temperature. Embryos were washed multiple times with 0.1% saponin/1X PBS before imaging. Antibody information is provided in **Supplementary File 1**.

IHC for quantification of cardiomyocytes in the AVC required the use of two rabbit primary antibodies (anti-DsRed and anti-Vmhc (Song et al., 2019)). To accommodate this, after blocking, embryos were first incubated with anti-Amhc (anti-Myh6/S46, mouse IgG1) and anti-DsRed (rabbit) overnight at 4⁰C. Secondary antibodies (anti-mouse IgG1-Dylight 405 and anti-rabbit-TRITC) were then applied for 2 hours at room temperature. After washing multiple times with 0.1% saponin/1X PBS, embryos were again incubated in saponin blocking solution for 1 hour. Anti-Vmhc (rabbit) was then diluted in saponin blocking solution and applied to embryos overnight at 4⁰C. Embryos were then washed multiple times with 0.1% saponin/1X PBS. Anti-rabbit-Alexa-488 was diluted in saponin blocking solution and embryos were incubated in secondary antibody for 1 hour at room temperature, followed by multiple washes with 0.1% saponin/1X PBS.

Following staining, embryos were imaged on a Nikon A1 inverted confocal microscope. Denoise-AI was employed on images taken using an HD resonance scanner. Cardiomyocytes were counted using Photoshop and ImageJ. For quantification of PCs, DsRed^+^/Amhc^+^/*fgf13a:*EGFP^+^ cardiomyocytes were counted as PC, while the DsRed^+^/Amhc^+^/*fgf13a:*EGFP^-^ cardiomyocytes were counted as atrium. For AVC counting, DsRed^+^/Amhc^+^/Vmhc^-^ cardiomyocytes were counted as atrium, DsRed^+^/Amhc^+^/Vmhc^+^ cardiomyocytes were counted as AVC, and DsRed^+^/Amhc^-^/Vmhc^+^ cardiomyocytes were counted as ventricle.

### Heart rate analysis

To analyze heart rate, embryos were anesthetized in 0.16 mg/mL tricaine and mounted in 1% low-melt agarose. Hearts were imaged on a Nikon Ti-2 SpectraX Widefield microscope with an Andor Xyla 4.2 megapixel, 16-bit sCMOS monochromatic camera at 200 frames per second (fps) in a controlled temperature chamber (28.5⁰C). Analysis performed in Nikon Elements software.

### Heat-shock experiments

Embryos resulting from a cross of adult *nr2f1a^+/-^*; *Et(fgf13a:EGFP); Tg(myl7:DsRed2-NLS)* and *nr2f1a^+/^; Tg(hsp70l:nkx2.5-EGFP)* zebrafish were first sorted for the *Et(fgf13a:EGFP)* transgene, as it is also expressed at low levels in the epidermis and in the olfactory pits (Poon et al., 2016). At 20 or 40 hpf, 10 *Et(fgf13a:EGFP)^+^* embryos per tube were placed into 0.5 ml PCR tubes with 200 ul embryo water and heat-shocked for 30 minutes by raising the temperature to 37⁰C using a thermocycler. Embryos were then returned to petri dishes and placed in a 28.5⁰C incubator. 3-4 hours after the heat-shock, when the Nkx2.5-EGFP becomes visible, embryos were sorted based on the presence of Nkx2.5-EGFP, which could be distinguished over the fluorescence from the *Et(fgf13a:EGFP)* transgene. Embryos were then returned to the 28.5⁰C incubator and allowed to develop until the desired timepoints when they were harvested, processed for IHC, imaged with confocal microscopy, and the cardiomyocytes counted. Embryos without the *Tg(hsp70l:Nkx2.5-EGFP)* transgene that were heat-shocked served as controls. All embryos were genotyped for the *nr2f1a* mutant allele.

### Fluorescence activated cell sorting (FACS)

GFP^+^ cardiomyocytes from wild-type and *nr2f1a* mutant with the *Tg(amhc:EGFP)*, *Tg(myl7:EGFP)*, or *Tg(amhc:EGFP)*; *Tg(myl7:DsRed2-NLS)* transgenes were isolated using FACS at 48 or 96 hpf, as indicated. Briefly, embryos were dissociated as described previously (Holowiecki et al., 2020; Samsa et al., 2016; Stachura & Traver, 2011) with the following modifications. WT and *nr2f1a* mutant embryos were harvested, rinsed in Hank’s Balanced Salt Solution (HBSS) with Ca^2+^ and Mg^2+^, and transferred to Eppendorf tubes in 500µL of FACS media (0.9x PBS, 2% fetal bovine serum). 60µL of Liberase (Roche) was added and a pipette was used to triturate the embryos. Embryos were then incubated at 32.5⁰C for 15 minutes before being triturated again. This process was repeated until the embryos were completely dissociated. Following dissociation, the cell suspensions were centrifuged at 1,300 rpm (100 x g) for 5 minutes at 4⁰C. The supernatant was discarded, and the pellets were resuspended in a total volume of 400µL. Samples were then filtered through a 40 µm strainer and collected into a FACS tube. FACS was performed by the CCHMC Research Flow Cytometry Core using a 70 µm nozzle and cells were collected into Eppendorf tubes for downstream applications.

### Real time quantitative PCR (RT-qPCR)

2229 WT and 704 *nr2f1a* mutant *Tg(myl7:EGFP)^+^* cardiomyocytes were captured at 96 hpf. PolyA RNA was collected from the cardiomyocytes using the Single Cell RNA Purification Kit (Norgen Biotek) and then amplified using the SeqPlex RNA Amplification Kit (Sigma-Aldrich), which generates double-stranded cDNA. RT-qPCR was performed using SYBR Green Mix in a BioRad CFX-96 PCR machine. Expression levels were standardized to β-actin and analyzed using the 2^−ΔΔCT^ Livak Method. Primer sequences are provided in **Supplementary File 2**.

### RNA-seq and ATAC-seq analysis

For RNA-seq, 7797 and 4118 *Tg(amhc:EGFP)^+^* cardiomyocytes, respectively, were captured from flow-sorting of ∼200 WT and *nr2f1a* mutant embryos at 48 hpf. RNA was collected using the Single Cell RNA Purification Kit (Norgen Biotek). PolyA RNA was then amplified using the MessageAmpII aRNA Amplification Kit (Thermo Fisher) and submitted to the CCHMC Sequencing core for quality control and library generation. Single-end sequencing was performed. RNA-seq data were submitted to the CCHMC Bioinformatics (BMI) core for analysis. To assess the need for trimming of adapter sequences and bad quality segments, the RNA-seq reads in FASTQ format were first subjected to quality control using FastQC v0.11.5, Trim Galore! V0.4.2, and Cutadapt v1.9.1 (Andrews, 2010; Krueger, 2012; Martin, 2011). The trimmed reads were aligned to the reference zebrafish genome version GRCz10/danRer10 with the program STAR v2.5.2 (Dobin et al., 2013). Aligned reads were stripped of duplicate reads using Picard v1.89 (Broad Institute, 2018). Gene-level expression was assessed by counting features for each gene, as defined in the NCBI’s RefSeq database (H. Li et al., 2009). Read counting was done with the program feature Counts v1.5.3 from the Rsubread package (Liao et al., 2019). Raw counts were normalized as transcripts per million (TPM). Differential gene expressions between conditions was assessed with the R v3.4.4 package DESeq2 v1.18.1 (Love et al., 2014).

For ATAC-seq, 8230 and 4338 *Tg(amhc:EGFP)^+^*; *Tg(myl7:DsRed2-NLS)^+^* cardiomyocytes, respectively, were captured from flow-sorting of ∼200 WT and *nr2f1a* mutant embryos at 48 hpf. ATAC-seq libraries were generated from the cells using the Nextera Kit (Illumina), as previously reported (Buenrostro et al., 2015). Libraries were submitted to the CCHMC for paired-end sequencing. ATAC-seq data were then submitted to the CCHMC BMI Core for analysis. Reads in FASTQ format were subjected to quality control, as above. The trimmed reads were aligned to the zebrafish genome (GRCz10/danRer10) using STAR v2.5.2 with splice awareness turned off. Aligned reads were stripped of duplicate reads using Picard v1.89. Peaks were called using MACS v2.1.2 using the broad peaks mode (Y. Zhang et al., 2008). Peaks with q-value <0.01 were selected for further analysis. Common peaks among all samples were obtained by merging called peaks at 50% overlap using BEDtools v2.27.0 (Quinlan & Hall, 2010). The common peaks, originally in BED format were converted to a Gene Transfer Format (GTF) to enable fast counting of reads under the peaks with the program feature Counts v1.5.3 (Rsubread package) (Liao et al., 2019). Differential open chromatin relative to control samples was assessed with the R package DESeq2 v1.18.1 (Love et al., 2014). Each differentially open chromatin region was annotated with closest differentially expressed gene from the RNA-seq. Differentially open chromatin regions are selected for motif enrichment analysis using HOMER v4.10 (Heinz et al., 2010) and known zebrafish motifs from CIS-BP v1.92 (Weirauch et al., 2014) are used to detect motif occurrences.

Accession numbers for RNA-seq and ATAC-seq data used for analysis will be deposited in GEO and supplied prior to acceptance of the manuscript.

### Electrophysiological analysis of hearts

Optical mapping of action potential duration and conduction velocities were performed similar to what was previously reported (Colombo et al., 2018; Mosimann et al., 2015; Panáková et al., 2010). Briefly, hearts from zebrafish embryos were isolated at 48, 72, and 96 hpf and stained with the transmembrane potential-sensitive dye FluoVolt^TM^ (Invitrogen) for 20’. To measure action potential duration, fluorescence intensities were recorded with a high-speed CCD camera (Redshirt Imaging) with 14-bit resolution at 2000 frames/second. Action potentials were temporally (800 Hz low-pass cutoff) and spatially (3×3 pixel average) filtered to enhance signal-to-noise ratios. Action potential duration at 20% repolarization (APD20) was defined as the absolute time difference between the decay phase and the upstroke phase of the local action potential at 20% repolarization and was measured at the midpoint of the atria. Conduction velocity vectors across the hearts were derived using an established algorithm (Bayly et al., 1998) with custom scripts in MATLAB (Version R2018B, Mathworks) and estimated from the local action potentials (Panáková et al., 2010). All electrophysiological measurements were averaged across the relevant anatomical segments for each heart. Embryos were genotyped following analysis.

### Statistical analysis

Comparisons between groups were analyzed using one- and two-way ANOVAs with multiple comparisons. Fisher’s exact test was used to determine if two proportions were statistically distinct. Statistical analysis was performed using GraphPad Prism. A p-value of <0.05 was considered statistically significant for all analysis. Error bars indicate s.e.m.

## Acknowledgements

We thank members of the Waxman lab for fruitful discussions and the CCHMC veterinary staff for care of the zebrafish. This work was supported by funding from the National Institutes of Health: [R01 HL137766, R01 HL141186, and R01 HL154522 to J.S.W.], [F31 HL152600 to K.E.M], and [R24 OD017870 and the Leducq Foundation to C.A.M.]

## Supplementary Information

**Figure 1 – figure supplement 1.**
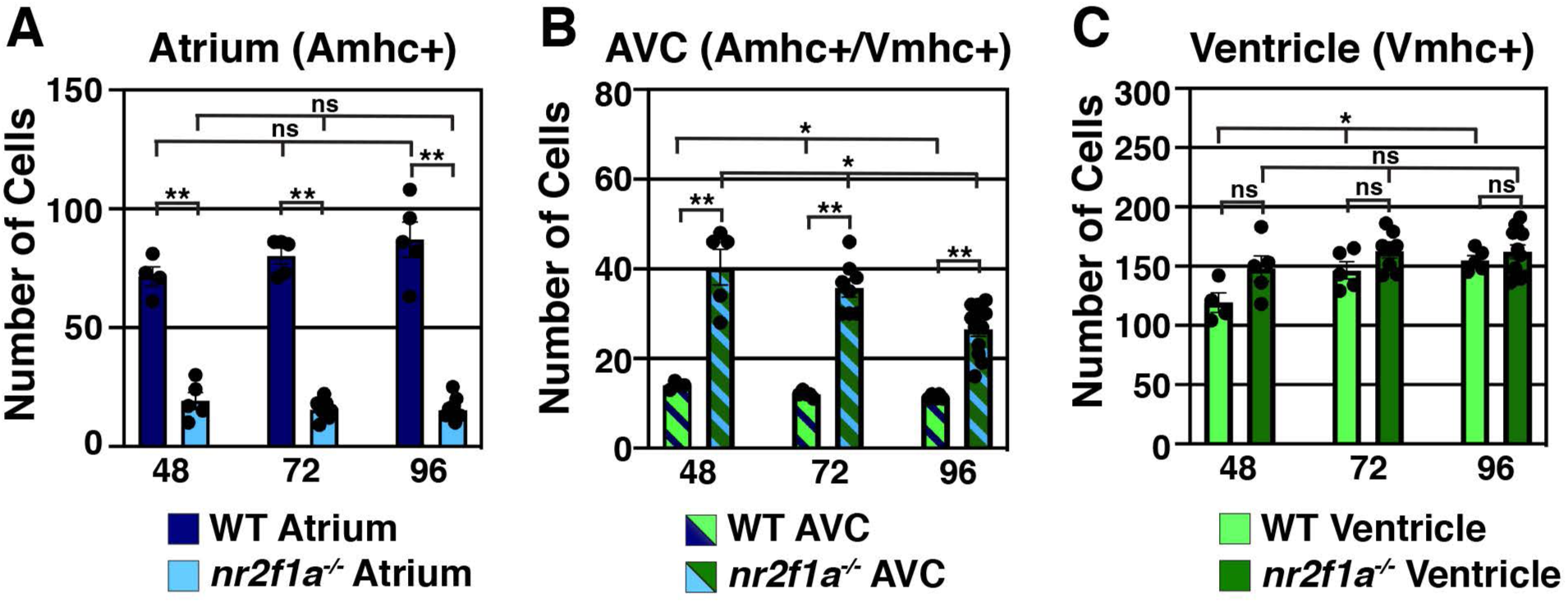
Quantification of cardiomyocytes in *nr2f1a* mutants. **A)** Quantification of ACs (Amhc^+^ only). **B)** Quantification of AVC cardiomyocytes (Amhc^+^/Vmhc^+^). **C)** Quantification of VCs (Vmhc^+^ only) cardiomyocytes. 48 hpf: WT (n = 4), *nr2f1a*^-/-^ (n = 5); 72 hpf: WT (n = 5), *nr2f1a*^-/-^ (n = 8); 96 hpf: WT (n = 5), *nr2f1a*^-/-^ (n = 12). Graphs are the same data presented in Figure 1J,K,L. Differences between WT and *nr2f1a^-/-^* were analyzed using ANOVA with multiple comparisons. * p = 0.05 – 0.001, ** p < 0.001

**Figure 2 – figure supplement 1.**
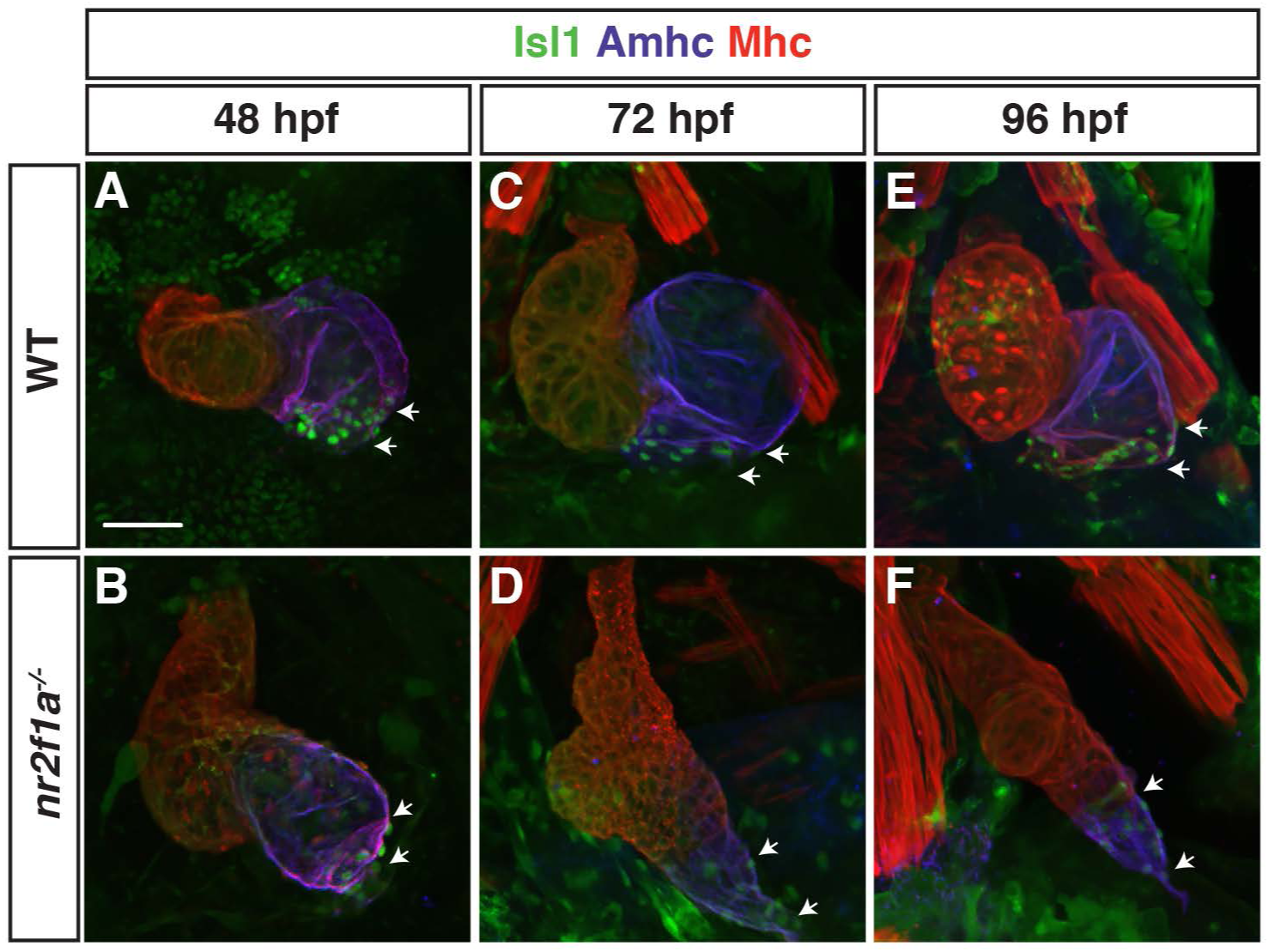
Isl1 expression expands throughout the atrium in *nr2f1a* mutant embryos. **A-F)** Mhc (red), Amhc (purple) and Isl1 (green) expression in WT and *nr2f1a* mutant embryos at 48, 72, and 96 hpf. White arrows mark boundaries of Isl1 expression at the venous pole. Number of embryos examined: 48 hpf: WT (n = 5), *nr2f1a^-/-^* (n = 5); 72 hpf: WT (n = 5), *nr2f1a^-/-^* (n = 5); 96 hpf: WT (n = 6), *nr2f1a^-/-^* (n = 10). Scale bars indicate 50 µm.

**Figure 3 – figure supplement 1.**
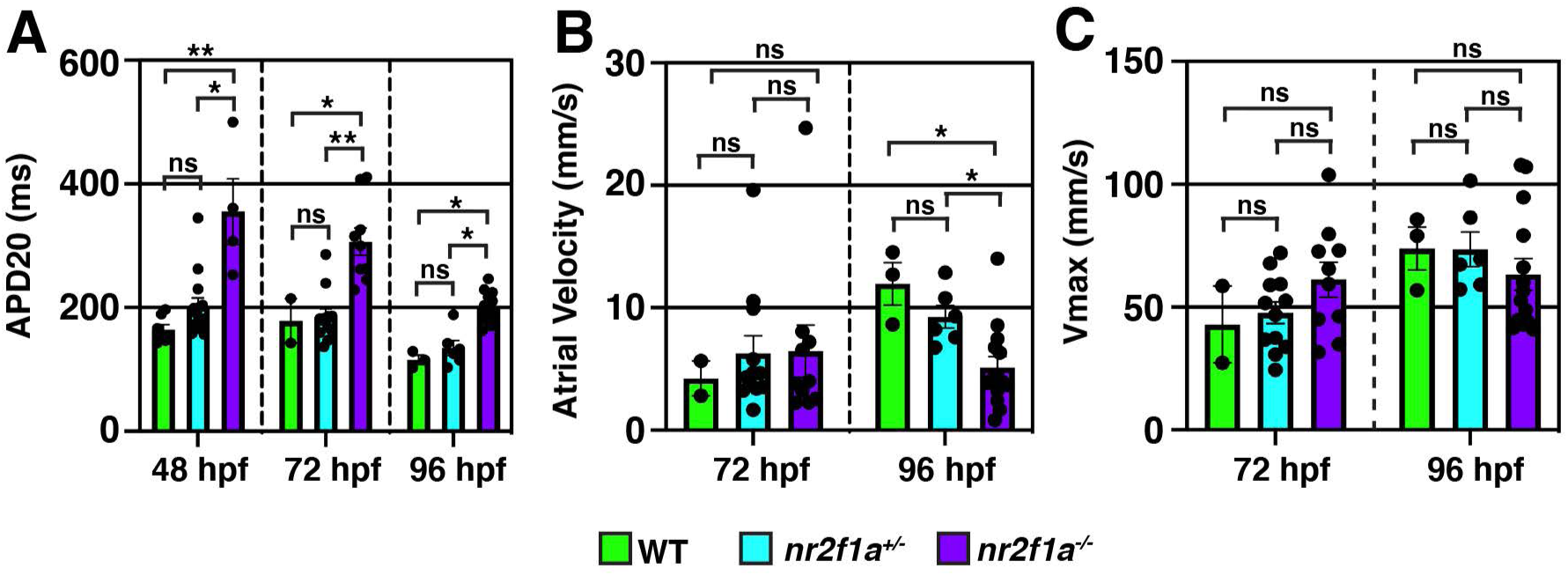
Prolonged repolarization and decreased atrial conduction velocity in *nr2f1a* mutants compared to WT and heterozygous *nr2f1a* atria. A) APD20 of WT, *nr2f1a^+/-^*, and *nr2f1a^-/-^* hearts. **B)** Atrial velocity of WT, *nr2f1a^+/-^*, and *nr2f1a^-/-^* hearts. **C)** Vmax of WT, *nr2f1a^+/-^*, and *nr2f1a^-/-^* hearts. Error bars indicate s.e.m. 48hpf: WT (n = 7), *nr2f1a^+/-^* (n = 14), *nr2f1a^-/-^* (n = 4); 72 hpf: WT (n = 2), *nr2f1a^+/-^* (n = 11), *nr2f1a^-/-^* (n = 9); 96 hpf: WT (n = 3), *nr2f1a^+/-^* (n = 6), *nr2f1a^-/-^* (n = 14). Differences between WT and *nr2f1a^-/-^* were analyzed using ANOVA with multiple comparisons. * p = 0.05 – 0.001, ** p < 0.001

**Figure 4 – figure supplement 1.**
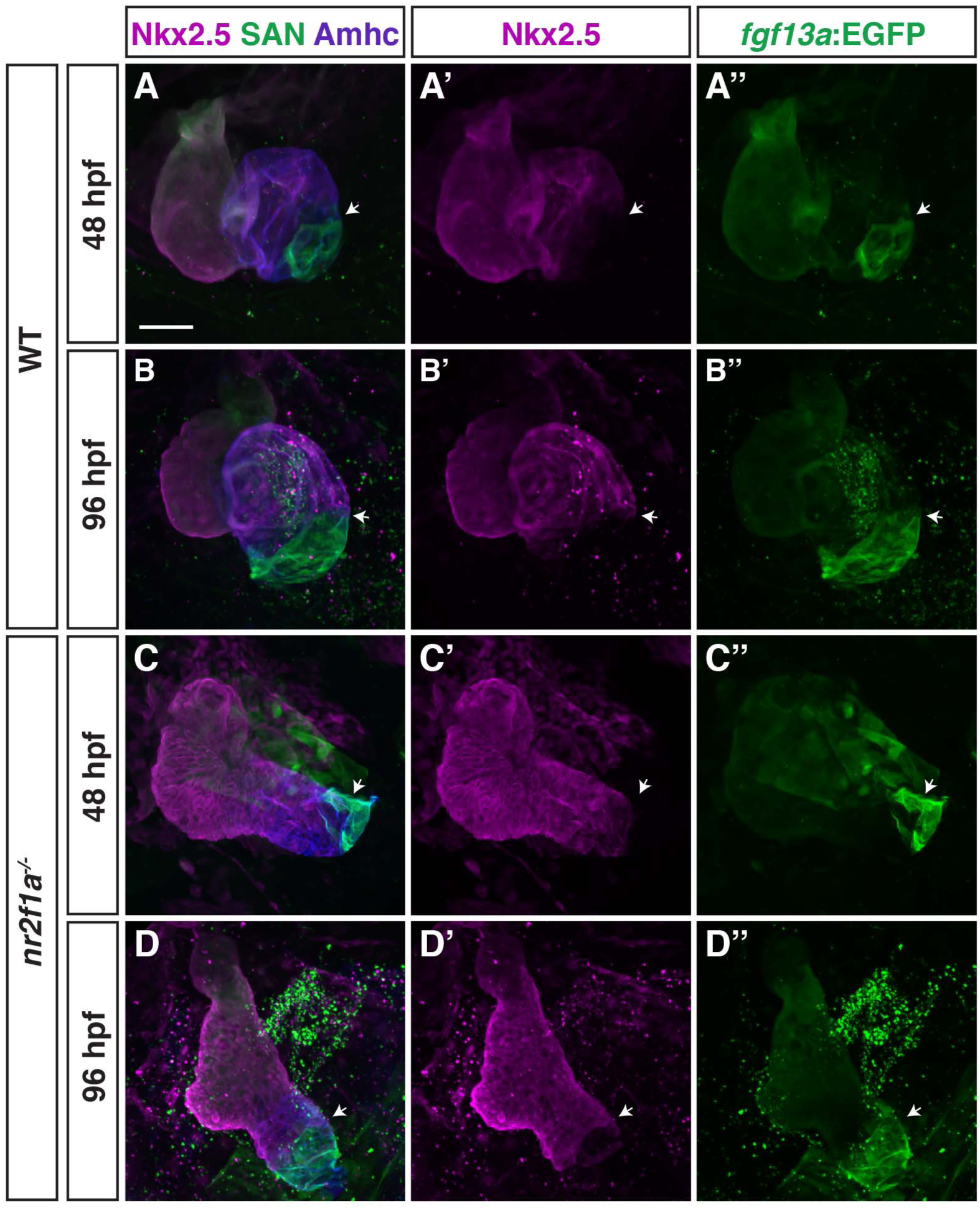
*nkx2.5*:ZsYellow expression within the atrium recedes toward the arterial pole in *nr2f1a* mutants. **A-D”)** IHC for *nkx2.5*:ZsYellow (purple), Amhc (blue), and *fgf13a*:EGFP (green) in the atria of WT and *nr2f1a* mutant embryos. White arrows indicate border of *nkx2.5*:ZsYellow and *fgf13a*:EGFP expression within the atria. Number of embryos examined at 48 hpf: WT (n = 8), nr2f1a-/- (n = 10); 96 hpf: WT (n = 9), nr2f1a-/- (n = 12). Scale bar indicate 50 µm.

**Figure 5 – figure supplement 1.**
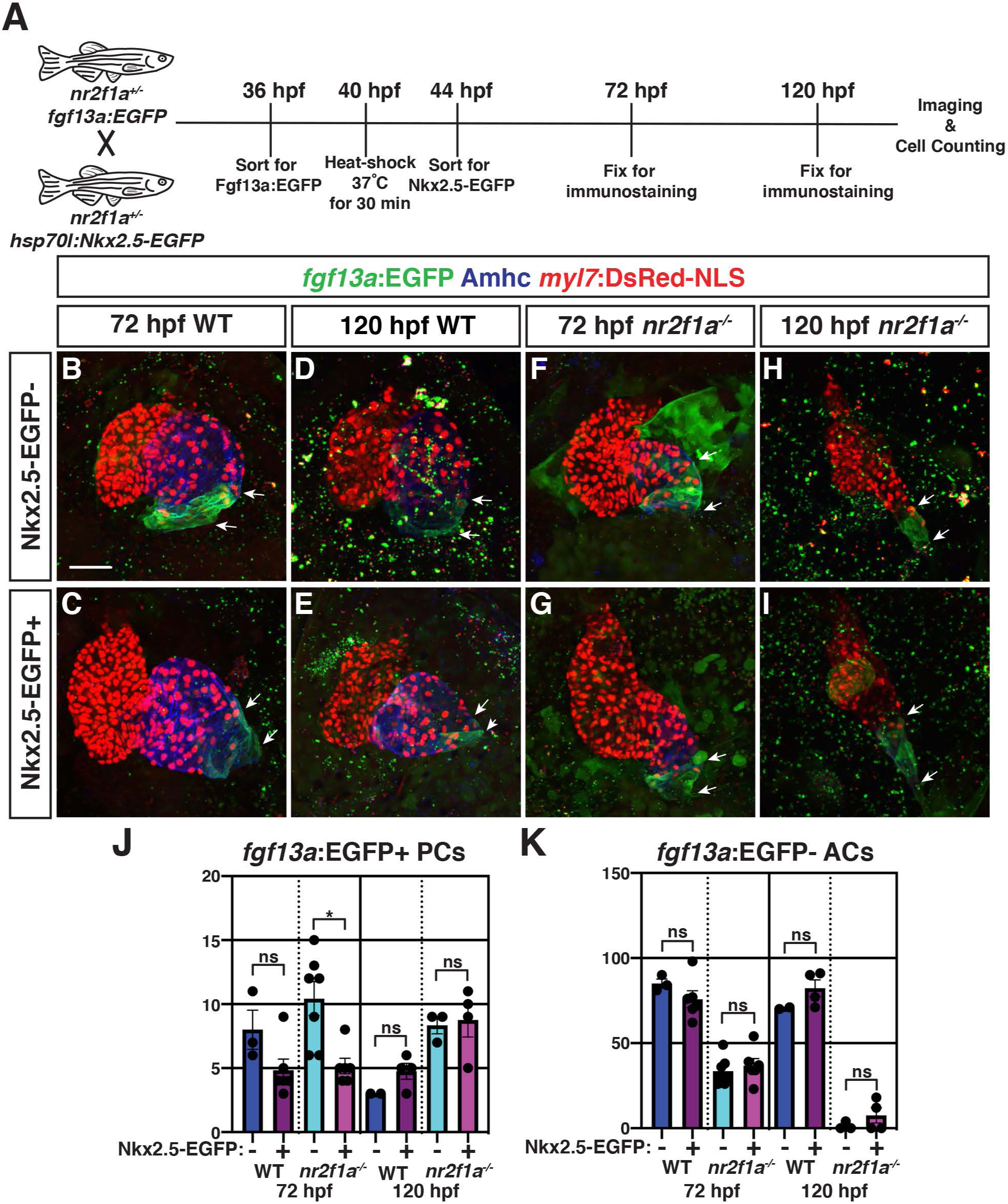
Nkx2.5-EGFP induction at 40 hpf partially represses the expansion of PC identity. **A)** Schematic of heat-shock experiments. **B-I)** IHC for *myl7*:DsRed-NLS (red), Amhc (blue), and *fgf13a*:EGFP (green) of representative hearts from WT and *nr2f1a* mutant embryos with and without Nkx2.5-EGFP. White arrows indicate boundaries of *fgf13a*:EGFP expression. **J-K)** Quantification of *myl7*:DsRed2-NLS^+^/Amhc^+^/*fgf13a*:EGFP^+^ (PCs) and *myl7*:DsRed2-NLS^+^/Amhc^+^/*fgf13a*:EGFP- (ACs) with and without Nkx2.5-EGFP induction at 72 and 120 hpf. Error bars indicate s.e.m. 72 hpf WT: Nkx2.5-EGFP^-^ (n = 3), Nkx2.5-EGFP^+^ (n = 6); 72 hpf *nr2f1a^-/-^*: Nkx2.5-EGFP^-^ (n = 7), Nkx2.5-EGFP^+^ (n = 6); 120 hpf WT: Nkx2.5-EGFP^-^ (n = 2), Nkx2.5-EGFP^+^ (n = 4); 120 hpf *nr2f1a^-/-^*: Nkx2.5-EGFP^-^ (n = 3), Nkx2.5-EGFP^+^ (n = 4). Scale bars indicate 50 µm. Differences between WT and *nr2f1a^-/-^* were analyzed using ANOVA with multiple comparisons. * p = 0.05 – 0.001, ** p < 0.001

**Figure 6 – figure supplement 1.**
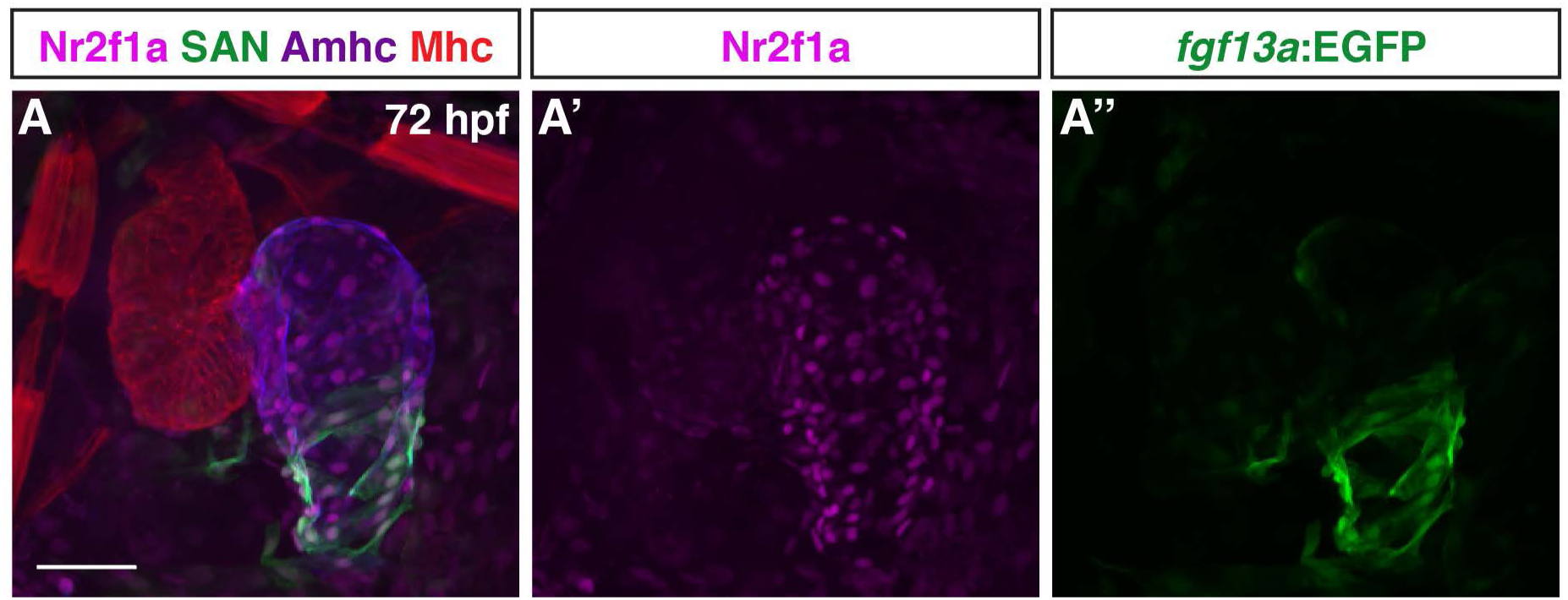
Nr2f1a is expressed throughout ACs in the atrium. **A-A”)** IHC for Nr2f1a (purple), *fgf13a*:EGFP (green), Amhc (blue), and Mhc (red) in WT hearts at 72 hpf (n = 7). Scale bar indicates 50 µm.

**Figure 1 – Source Data 1. (xlsx) RNA-seq and ATAC-seq data with differential gene expression, changes in called peaks, and putative Nr2f binding sites.**

**Figure 3 – Video 1. Heart from 48 hpf WT embryo.**

**Figure 3 – Video 2. Heart from 48 hpf *nr2f1a* mutant embryo.**

**Figure 3 – Video 3. Heart from 72 hpf WT embryo.**

**Figure 3 – Video 4. Heart from 72 hpf *nr2f1a* mutant embryo.**

**Figure 3 – Video 5. Heart from 96 hpf WT embryo.**

**Figure 3 – Video 6. Heart from 96 hpf *nr2f1a* mutant embryo.**

**Supplementary File 1.**
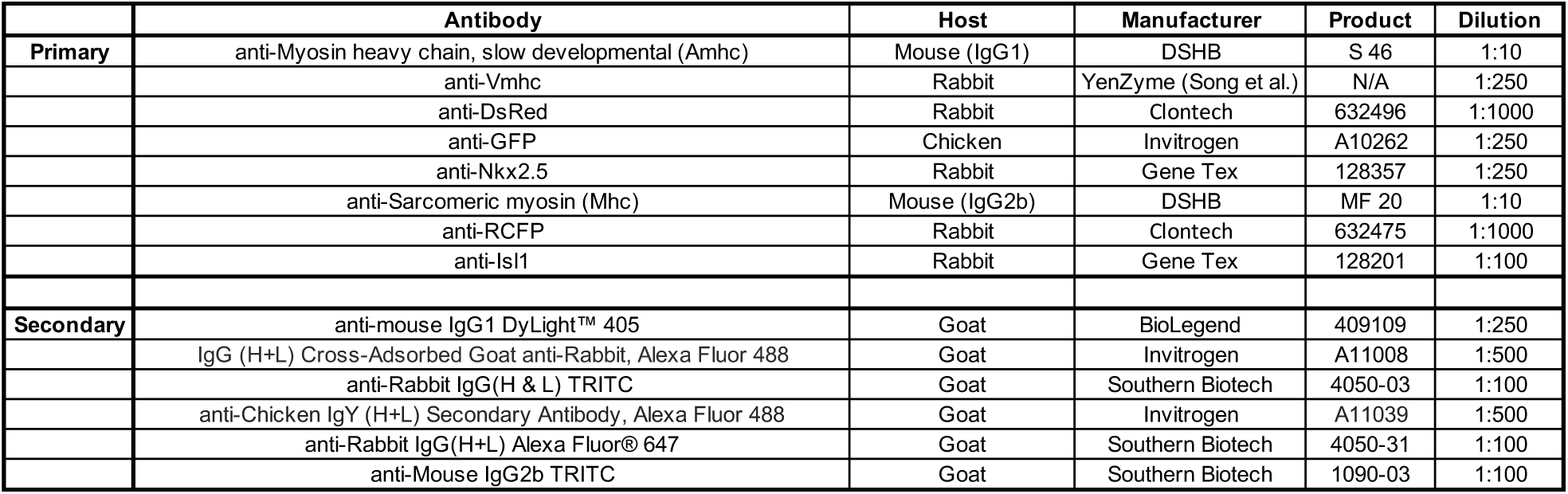
Antibody information.

**Supplementary File 2.**
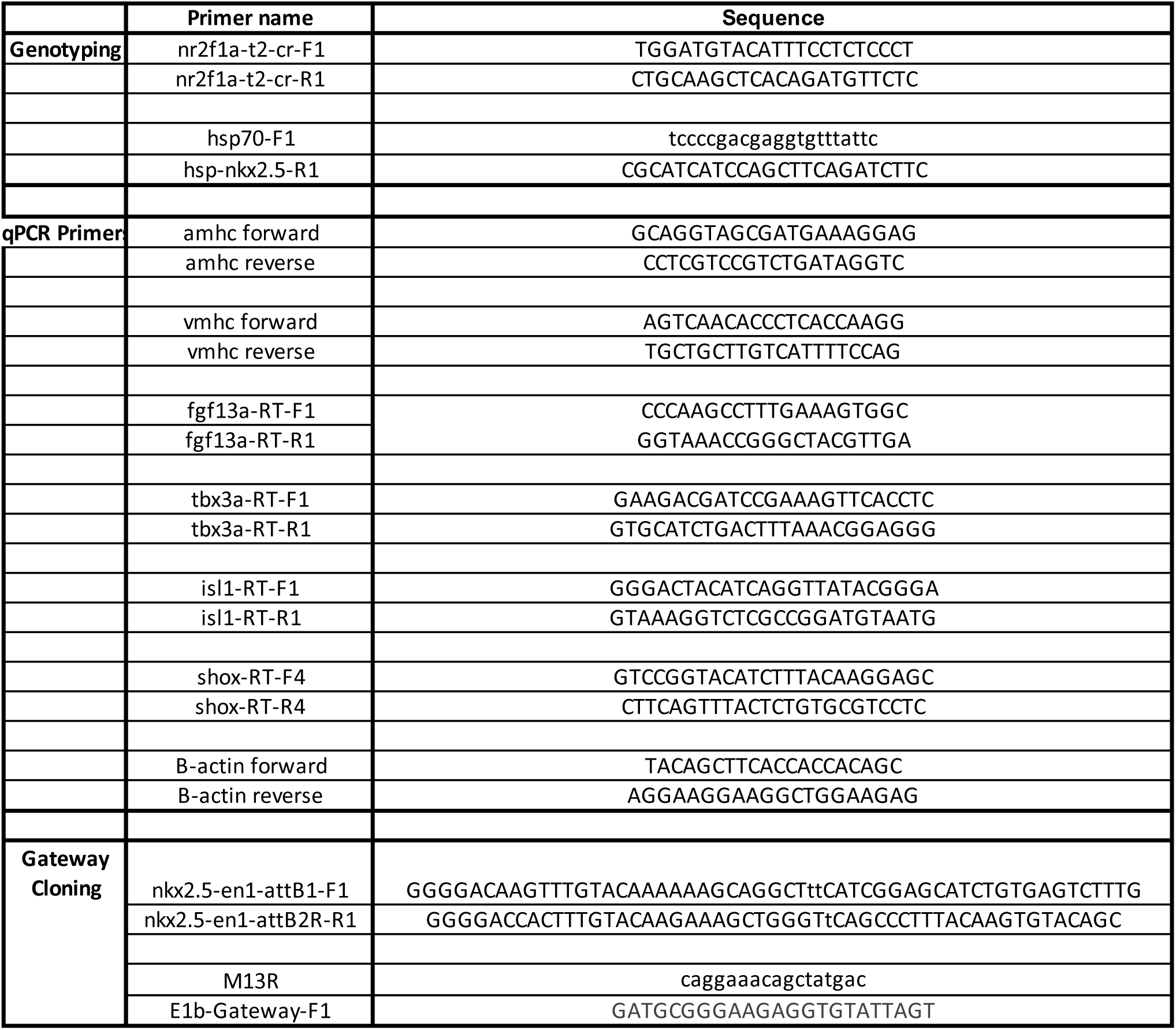
Primer information.

